# MCseg: reference-guided AI-agent development of a fixed cell segmentation and transcript attribution workflow for spatial transcriptomics

**DOI:** 10.64898/2026.09.20.752837

**Authors:** Chi-Ru Chan, Nai-Wen Chang, Chia-Yi Wang, Hsin-Yuan Tan, Sung-Jan Lin

## Abstract

Cell-level analysis of high-resolution spatial transcriptomics depends on how tissue images are segmented and how transcripts are assigned to cells, yet paired reference measurements for developing and evaluating such workflows are scarce and cannot be assumed for each sample requiring analysis. We developed MCseg using a single paired tissue section as a development reference: an AI agent iteratively composed and scored combinations of pretrained Cellpose models and image-processing operations against Xenium-derived cell boundaries, and the retained workflow was then fixed so that it can be deployed without the reference assay or the agent. In the lung adenocarcinoma development set, the fixed workflow increased mean panoptic quality from 0.432 to 0.472 relative to an Optuna-tuned two-diameter Cellpose baseline, with improvement in four of six regions. In an expert-annotated colorectal cancer region not used during development, MCseg showed higher lineage recall and micro-F1 (0.805 versus 0.732) than the StarDist-based ENACT workflow among 10,275 reference cells covered by both methods. Across 15 regions from one colorectal cancer section, MCseg showed higher neighborhood expression discordance and lower lineage-exclusive co-expression than Space Ranger at similar UMI density, while capturing fewer transcripts. The fixed workflow could also be applied to fresh-frozen breast cancer, a distinct tissue and preservation condition, without a new architecture search. These results show how a limited paired reference can be used to develop a fixed, reusable workflow that no longer depends on the reference assay; MCseg distributes this workflow as a locally run, no-code platform.

**Availability and implementation:** MCseg is available under the MIT License at https://github.com/ddmanyes/MCseg. The repository contains the segmentation workflow, transcript-attribution scripts, analysis code, AI-agent prompts, and development logs.

**Bullet points:**

1. A paired spatial reference AI-agent search for a fixed segmentation workflow
2. MCseg improves agreement with Xenium boundaries over a tuned Cellpose baseline
3. MCseg reduces lineage mixing relative to Space Ranger in colorectal cancer
4. A local no-code tool runs the fixed workflow without the reference assay or an AI agent

## 1. Introduction

Spatial transcriptomics links molecular profiles to tissue architecture and is increasingly used to resolve cellular heterogeneity and local tissue organization (Moses and Pachter, 2022; Stahl, et al., 2016). Compared to prior low-resolution methods, high-resolution platforms now approach cellular dimensions by different routes. Imaging-based systems such as Xenium detect individual RNA molecules and infer cell boundaries computationally, whereas sequencing-based platforms such as Visium HD measure the whole transcriptome on a dense array of 2-µm capture bins (Janesick, et al., 2023). The latter are subcellular sampling units rather than biological cells. Cell-level analysis therefore requires both a credible segmentation of the tissue image and an explicit rule for assigning spatially indexed expression measurements to those boundaries.

Cell segmentation is key to correctly assigning transcripts to corresponding cells. Errors in segmentation propagate directly into downstream biological analysis, compromising the correctness of the following bioinformatic analysis. For example, overexpanded masks can cross into neighboring cells, glandular lumina, or stromal spaces and mix transcripts from different lineages. Conversely, conservative nucleus-restricted masks reduce contamination but leave substantial cytoplasmic signal unassigned. Aggregating Visium HD data into 8- or 16-µm bins avoids explicit segmentation, but a single bin can span more than one cell and dilute rare-cell signals in densely packed regions (Long, et al., 2025; Petukhov, et al., 2022). Thus, segmentation is not simply an image-processing problem. It defines the unit on which cell states, neighborhoods, and interactions are inferred for downstream analysis.

Several general-purpose and spatially informed methods addressing this problem are now available (Blampey, et al., 2024; Chen, et al., 2023; Kamel, et al., 2025; Petukhov, et al., 2022; Polanski, et al., 2024), but no single strategy is uniformly optimal across tissue morphologies, staining conditions, and cell densities. Pretrained generalist segmentation models have reduced the need for task-specific training, yet applying them to a new tissue still requires choosing and combining preprocessing, models, mask integration, and boundary expansion. Standardized workflows are convenient but can overexpand boundaries in heterogeneous tissue, whereas more configurable approaches often require command-line tools, environment management, and repeated parameter tuning. These practical demands remain a barrier for groups that generate or interpret spatial transcriptomic data without dedicated image-analysis expertise.

Benchmarking is equally challenging. Expert-drawn whole-cell boundaries are expensive to generate and difficult to register exactly across assays (Greenwald, et al., 2022). Geometric scores are informative when matched reference boundaries are available, but they do not directly measure whether expression has been attributed to the correct biological cell. Transcript capture alone is also insufficient because a larger mask can collect more transcripts while increasing cross-cell contamination. A useful evaluation therefore needs to consider geometric fidelity, transcript recovery, mask-normalized signal density, and transcriptomic purity together (Jones, et al., 2025; Mitchel, et al., 2026; Plummer, et al., 2026). The same reference bottleneck affects workflow development as well as evaluation. Whole-cell boundaries are difficult to define with certainty from H&E images alone. Matched multimodal measurements, such as imaging-based spatial profiling of the same section, provide an opportunity to evaluate segmentation workflows against complementary spatial information, but cannot be assumed to be available for each sample requiring analysis. This motivates separating reference-guided workflow development from routine deployment

We therefore asked whether a paired tissue section could guide the composition of pretrained segmentation models and image-processing operations into a fixed workflow for subsequent analysis without the reference assay. Conventional hyperparameter optimization can improve a fixed researcher-specified pipeline, but it does not readily explore changes in model composition, preprocessing, mask integration, or boundary expansion. AI agents are increasingly being considered as tools for planning and iteratively refining bioinformatic workflows (Corpas, et al., 2026). We adapted the closed-loop logic of AutoResearch (Karpathy, 2025) to a narrower development task: an AI agent modified executable segmentation code within a researcher-defined library, evaluated each candidate against reference boundaries, and used the resulting score to propose the next workflow. The agent was used only during development. The final routine MCseg analysis uses the fixed, retained architecture and does not require an external AI service.

Here we present MCseg (Multiple Cellpose Segmentation), a downloadable no-code platform for cell segmentation and transcript attribution in high-resolution spatial transcriptomics. We first established a conventionally optimized two-diameter Cellpose workflow (2Cseg) as a baseline, then allowed the AI agent to search a broader space of Cellpose models and image-processing operations using co-registered Xenium boundaries from lung adenocarcinoma as the development reference. Each dataset was assigned a distinct role. The LUAD section supported workflow development and development-set geometric evaluation; an expert-annotated colorectal cancer (CRC) region and 15 regions from the same CRC section supported external lineage-assignment and transcript-derived evaluation; and a fresh-frozen breast cancer dataset tested whether the fixed workflow could be applied to a distinct tissue and preservation condition.

## 2. Materials and Methods

### 2.1 Study design and spatial transcriptomics datasets

Three publicly available datasets were used. The development dataset consisted of co-registered Visium HD and Xenium Prime data generated from the same formalin-fixed, paraffin-embedded lung adenocarcinoma (**LUAD**) section. Six fixed-size regions of interest (**ROI**s) were used for geometric evaluation, with Xenium cell boundaries serving as the reference (see **Supplementary Figure S1** for ROI locations). These registered data were also used during workflow development; consequently, all LUAD geometric results are reported as development-set analyses.

A colorectal cancer (**CRC**) Visium HD FFPE dataset was obtained from the Gene Expression Omnibus (GSE280318). Fifteen ROIs were used for transcript-derived benchmarking. These ROIs are non-overlapping regions of a single tissue section and were therefore treated as within-section replicates rather than independent biological specimens. The expert-annotated ENACT dataset (Kamel, et al., 2025), comprising 20,991 reviewed cell centroids from a non-overlapping region of the same public section, was used for external geometric and cell-type validation.

A fresh-frozen breast cancer Visium HD whole-slide dataset from the 10x Genomics dataset portal was used to examine cross-tissue transfer. After development was complete, the converged MCseg workflow was applied to the CRC and breast cancer data without a new architecture search. Dataset details and selected tissue-region coordinates are provided in **Supplementary Note 3** and **Supplementary Table S2**.

### 2.2 Segmentation methods

The primary comparisons included Space Ranger version 4 (SR), MCseg, the 2Cseg baseline, and nucleus-restricted segmentation (NUC). StarDist and Proseg were included as additional comparators in the CRC analyses. The ENACT workflow, which combines StarDist with weighted bin-area assignment, was evaluated in the expert-annotated CRC benchmark. Software versions and method-specific settings are listed in **Supplementary Note 2**.

### 2.3 Development of the 2Cseg baseline

2Cseg was defined as a two-diameter Cellpose workflow. Its parameters were optimized with Optuna over 50 trials on three LUAD ROIs, using mean panoptic quality (PQ) against Xenium reference boundaries as the objective. The selected configuration was then evaluated across the six LUAD ROIs and achieved a mean PQ of 0.432. This researcher-defined architecture served as the baseline for testing whether changes in workflow composition could provide additional improvement. Full search settings are provided in **Supplementary Note 2D**.

### 2.4 AI-agent optimization and the MCseg workflow

The evaluation loop of AutoResearch (Karpathy, 2025), originally developed for language-model experiments, was adapted for H&E-based cell segmentation. The objective function was replaced with average precision at an intersection-over-union threshold of 0.5 (AP@0.5), calculated against the Xenium reference boundaries.

The agent was given a constrained library containing Cellpose cyto2, cyto3, nuclei, and cpsam models (Kirillov, et al., 2023; Stringer and Pachitariu, 2025; Stringer, et al., 2021), together with contrast-limited adaptive histogram equalization (CLAHE), HED stain unmixing, watershed segmentation, and Voronoi tessellation. Within a sandboxed environment, it modified a single Python script, ran the candidate workflow, recorded the score, and used the best-performing code, experimental log, and search memory to propose the next candidate. Each iteration was limited to 300 s.

Candidate workflows were generated and evaluated iteratively until further gains became small. The researchers defined the tool library, reference data, scoring function, prompt, and execution constraints, and reviewed the retained configuration before deployment. The prompts, execution scripts, and search records are available in the MCseg repository.

The converged workflow comprised CLAHE-based preprocessing, seven Cellpose passes (four cyto3 passes using multiple inputs and diameter settings and three cpsam passes), priority-based mask integration, optional transcript-density rescue, and Voronoi-constrained expansion. The full algorithm specification and step-by-step pipeline are provided in Supplementary Note 1. The deployed MCseg platform runs this fixed architecture; end users do not repeat the AI-agent search.

### 2.5 ENACT benchmarking

The ENACT expert-annotated CRC dataset (Kamel, et al., 2025) was used as an external benchmark. The ENACT reference centroids were initially detected with StarDist (Schmidt, et al., 2018) and then reviewed manually. This design may favor nuclear-centric methods in coverage analyses and was considered when interpreting the results.

A reference centroid located within an MCseg mask was counted as covered. Expression profiles were classified with CellTypist version 2 and mapped to the epithelial, stromal, and immune categories used by ENACT. Precision, recall, and micro-averaged F1 were calculated. To make each denominator explicit, we report the number of reference cells at each step: all reference centroids (n = 20,991), centroids covered by MCseg, covered cells with CellTypist labels mapped to the ENACT categories (n = 11,709), and cells covered by both MCseg and the ENACT workflow (n = 10,275). Subset analyses were restricted to cells covered by both MCseg and the ENACT workflow, whereas the overall analysis treated uncovered reference cells as false negatives.

Coverage efficiency was defined as the fraction of reference centroids covered divided by fractional transcript capture (FTC). For the ENACT comparator, weighted bin-area assignment used a 7 × 7-pixel sampling window around each bin centroid, with UMI counts distributed according to the effective overlap with neighboring masks (Kamel, et al., 2025).

### 2.6 Geometric evaluation

Geometric performance was measured with AP@0.5 and PQ. A predicted cell and a reference cell were matched when their intersection over union was at least 0.5. AP@0.5 summarizes detection precision and recall at this threshold.

PQ was calculated as segmentation quality (SQ) multiplied by recognition quality (RQ). SQ is the mean intersection over union of matched pairs, and RQ is the F1 score based on matched cells, false-positive predictions, and unmatched reference cells. PQ therefore captures both boundary conformity and detection completeness. Two MCseg evaluation settings were distinguished. The fixed-parameter deployment analysis used the repository’s default expansion setting and did not access the reference masks during segmentation. A separate reference-guided calibration analysis used the Xenium masks to select, independently for each ROI, the best-performing single-pass starting mask and the expansion strategy and distance (Supplementary Note 1, Steps 3 and 5). This analysis describes performance under sample-specific calibration with access to the reference; it is not a strict upper bound and was not treated as routine deployment performance.

### 2.7 Transcript attribution and RNA quality metrics

Each in-tissue 2-µm bin was assigned to the segmentation mask containing its full-resolution pixel centroid. Bin coordinates were converted to ROI-local pixel coordinates and used to retrieve the corresponding mask label. Bins with mask label 0 were treated as unassigned. Gene counts from bins sharing the same mask label were summed with sparse matrix operations to generate a cell-by-gene AnnData matrix. The same tissue region, defined by the Space Ranger tissue-detection mask, was used as the denominator for comparisons across methods.

Fractional transcript capture (FTC) was the proportion of tissue UMIs assigned to any cell mask. UMI density was the number of assigned UMIs divided by total mask area and was expressed as UMIs/µm^2^.

Neighborhood expression discordance (NED) measured transcriptomic separation between adjacent cells. For each ROI, the top 1,000 highly variable genes were selected, and per-cell counts were converted to relative frequencies. Adjacent masks were identified by grayscale dilation, and NED was calculated as the mean Hellinger distance between neighboring expression profiles. NED ranges from 0 to 1; higher values indicate greater separation between neighboring profiles. Because it is calculated within each segmentation geometry, NED is a reference-free measure of boundary sharpness rather than absolute geometric accuracy.

Lineage mixing was assessed with four marker pairs expected to be largely mutually exclusive: EPCAM–CD3E, MUC2–NKG7, ACTA2–CD3E, and PECAM1–EPCAM. A cell was counted as co-positive when both genes had at least one raw UMI. The lineage-exclusive co-expression rate was the proportion of cells meeting this criterion.

### 2.8 Clustering, spatial analysis, and statistics

Cell-level matrices were analyzed with Scanpy version 1.10 (Wolf, et al., 2018). Principal-component analysis used 30 components, the nearest-neighbor graph used 15 neighbors, and clusters were identified with the Leiden algorithm (Traag, et al., 2019). UMAP was calculated with min_dist = 0.5.

The CRC tertiary lymphoid structure (TLS) ROI was selected by local Moran’s I analysis of a composite score of MS4A1, CD79A, CXCL13, IGKC, CD3E, and LTB, using 8,999 permutations. Whole-slide distances among SPP1-positive macrophages, regulatory T cells (Tregs), and CD8-positive T cells were compared with complete spatial randomness distributions generated by permutation.

Paired ROI comparisons used Wilcoxon signed-rank tests or sign-flip permutation tests, as stated in the corresponding analyses. Sign-flip tests used 10,000 iterations, and 95% confidence intervals were obtained by bootstrap resampling. The Friedman test was used for comparisons involving several methods. Because the 15 CRC ROIs were drawn from a single tissue section, these tests describe consistency across regions within that section and do not support inference across patients. False-discovery-rate correction was applied where multiple tests were performed.

### 2.9 Software implementation

Analyses were performed in Python 3.10 or later with Cellpose, Scanpy, SciPy, and related image-processing libraries. The reported analyses were run on an Apple Silicon computer with Apple Metal Performance Shaders (MPS) acceleration. Runtime varied with ROI dimensions and cell density; the six geometric benchmark runs required approximately 20–40 min per ROI. MCseg also supports CPU-only execution and does not require a dedicated CUDA-enabled GPU, although CPU runtime was not benchmarked systematically in this study. MCseg is released under the MIT License.

## 3. Results

### 3.1 Iterative workflow search produced a multi-model MCseg segmentation engine

We first established the performance of the researcher-defined 2Cseg baseline. After Optuna optimization, 2Cseg reached a mean PQ of 0.432 ± 0.037 across the six LUAD ROIs, providing the reference for evaluating changes in workflow architecture (**Supplementary Figure S2**). We then opened workflow composition to an AutoResearch-inspired search. The agent evaluated combinations of preprocessing, Cellpose models, mask integration, and boundary expansion, ultimately retaining a workflow that combined CLAHE preprocessing, seven Cellpose passes across cyto3 and cpsam models at several diameter settings, priority-based mask integration with optional transcript-density rescue, and Voronoi-constrained expansion. On the primary development patch, AP@0.5 increased from 0.320 for the initial single-model workflow to 0.648 following the introduction of CLAHE preprocessing and Voronoi-constrained expansion, and further improved to 0.650 as the agent converged on the seven-pass multi-model architecture. Most of the improvement on this patch therefore preceded the seven-pass ensemble, which added a small further gain (**Supplementary Figure S11**). After researcher review, we froze this configuration as the deterministic segmentation engine used for subsequent analyses.

### 3.2 Fixed-parameter deployment and reference-guided calibration in the LUAD development set

We next characterized the frozen workflow under fixed-parameter deployment and compared it with both 2Cseg and a reference-guided calibration analysis. Without access to Xenium masks during segmentation, MCseg achieved a mean PQ of 0.472 ± 0.072 versus 0.432 ± 0.037 for 2Cseg, an absolute increase of 0.040 (9% relative). Here, geometric scores measure agreement with Xenium-derived reference boundaries, which are themselves computational estimates rather than error-free ground truth. This modest net change reflects a larger gain in segmentation quality (SQ) that was partially offset by a decrease in recognition quality (RQ), as detailed below; a substantially larger PQ gain (0.554, 28% relative) was obtained separately under reference-guided calibration. MCseg had the higher PQ in four of six ROIs (**Figure 2a-c; Supplementary Table S4)**. Because the LUAD section contributed to workflow development, these values should be interpreted as development-set performance rather than an independent generalization estimate.

**Figure 1.**
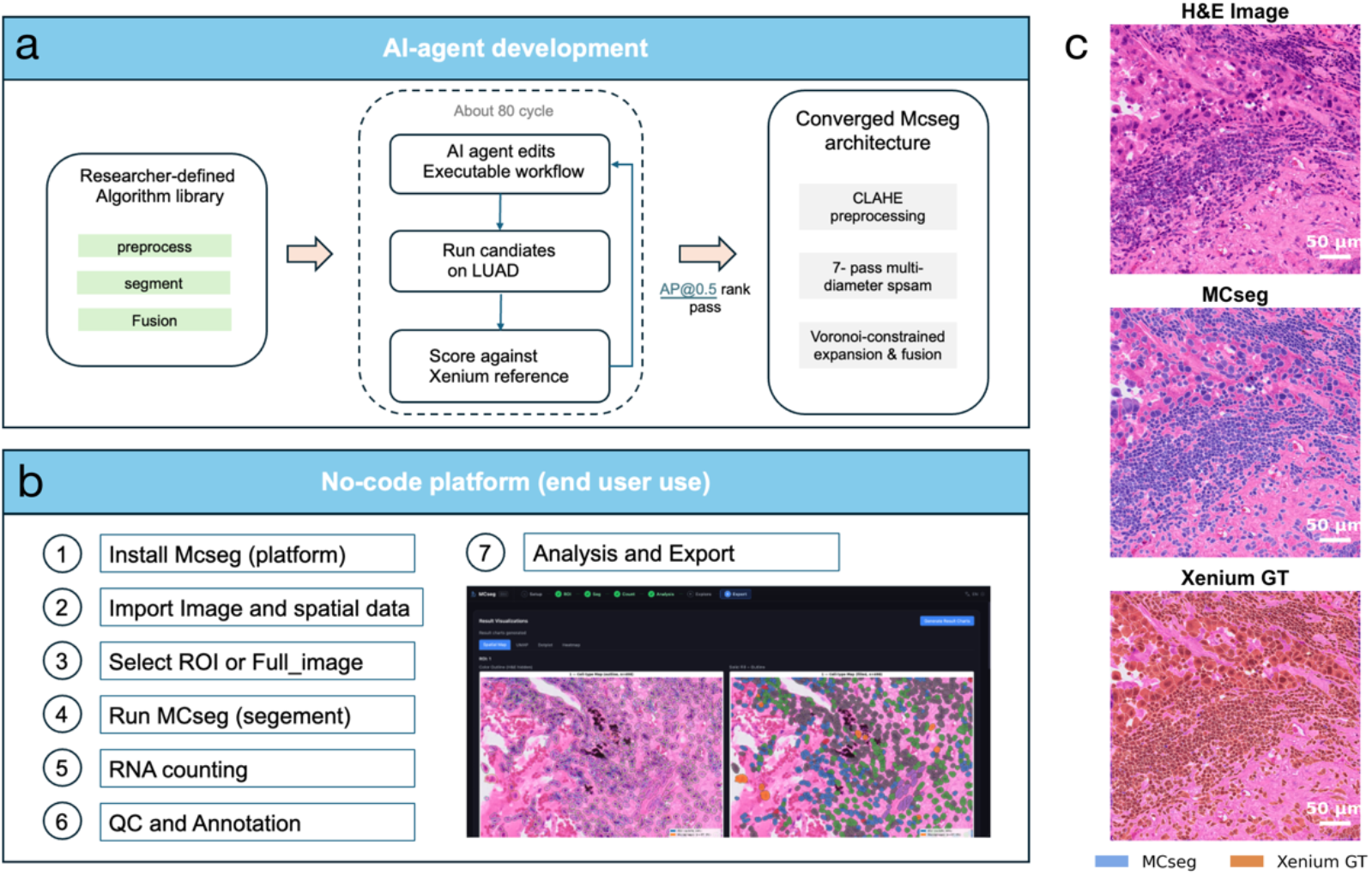
MCseg development and deployment. (a) Process of MCseg development. During development, an AI agent proposed and evaluated candidate segmentation workflows within a researcher-defined library and scoring framework. The retained components were reviewed and consolidated into a fixed seven-pass architecture. (b) Use platform. The deployed no-code platform accepts H&E and spatial expression data and returns cell masks and a cell-by-gene matrix without invoking the AI agent. (c) Representative images. The representative H&E image, MCseg masks, and Xenium reference boundaries are shown.

**Figure 2.**
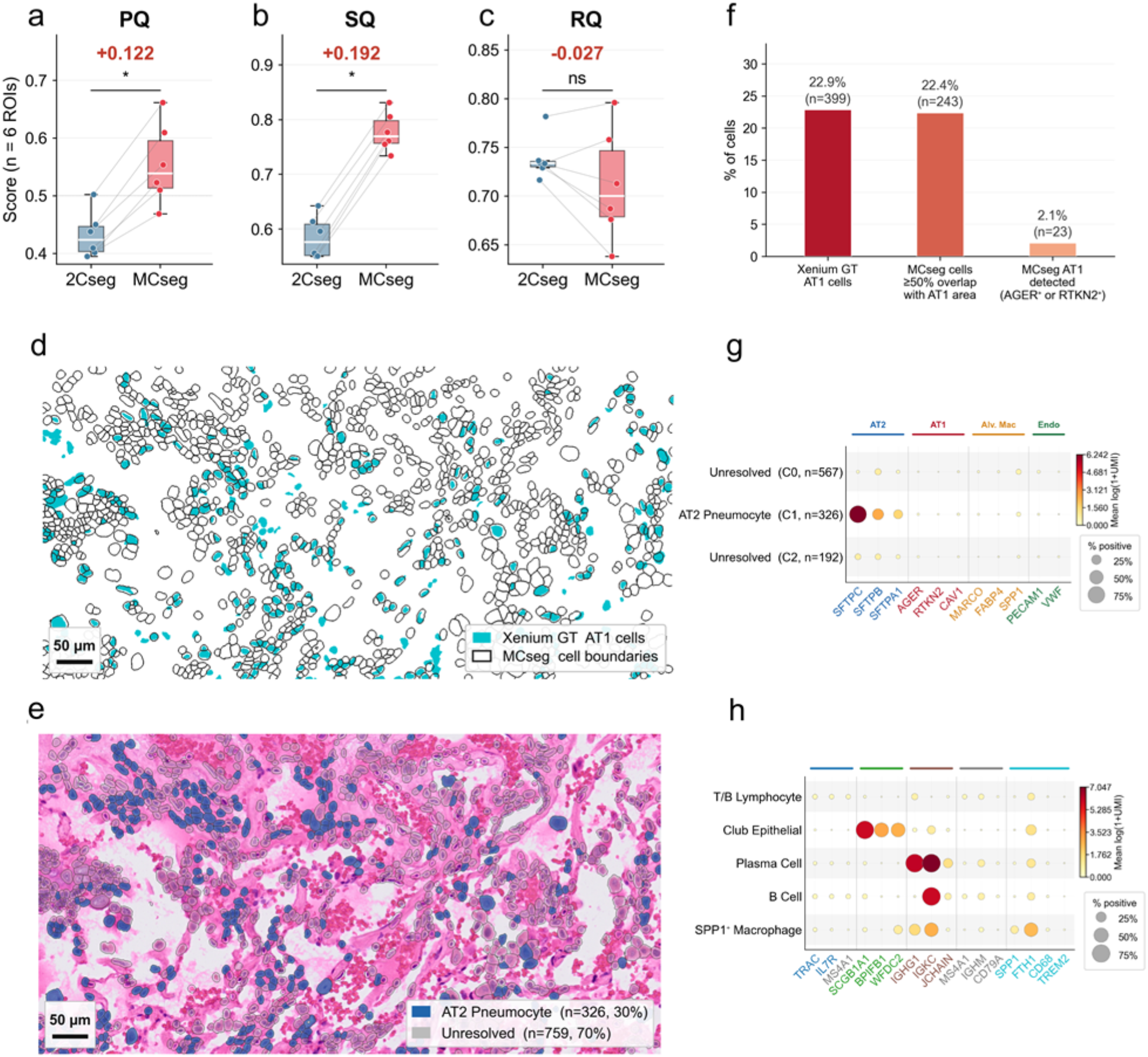
LUAD geometric benchmarking and biological evaluation. (a–c) Fixed-parameter deployment results for MCseg and 2Cseg across six LUAD ROIs using Xenium cell boundaries only for scoring. MCseg increased mean PQ and SQ, whereas RQ decreased; the reference-guided calibration analysis is reported separately in Supplementary Table S5. (d–f) Comparison of MCseg boundaries with Xenium-defined AT1 cells and the corresponding geometric and marker-based recovery rates. (g) Marker expression across LUAD cell clusters. (h) Cell-state marker expression in a macrophage-rich ROI. Scale bars are shown in the corresponding panels.

In the fixed deployment analysis, mean SQ increased from 0.585 to 0.736, but mean RQ decreased from 0.739 to 0.638, indicating a trade-off: matched boundaries conformed more closely to the reference, whereas some objects were missed or merged in dense regions. In the separate reference-guided calibration analysis, selecting the mask and the expansion strategy, and the distance independently for each ROI increased mean PQ to 0.554 ± 0.063, with SQ = 0.777 and RQ = 0.711 (**Supplementary Table S5; Supplementary Figure S3**). This calibration analysis exceeded 2Cseg in all six ROIs but used reference information to select the starting mask and expansion for each ROI; it therefore does not represent routine deployment and should not be read as a strict upper bound.

### 3.3 Cell-state analysis in challenging LUAD regions

We next examined two morphologically difficult regions: open alveoli containing thin alveolar type 1 (AT1) cells and tumor regions containing pigment-laden macrophages. In the alveolar region, MCseg assigned 22.4% of cells to boundaries overlapping Xenium-defined AT1 cells, close to the 22.9% AT1 fraction in the Xenium reference (**Figure 2d-f**). Direct marker-based recovery was much lower on Visium HD because only a small fraction of MCseg cells contained AGER or RTKN2 signal, consistent with the difficulty of sampling the very thin AT1 cytoplasm with 2-µm bins. AT2-associated cells, which contain more cytoplasm and RNA, formed a distinct cluster defined by SFTPC/SFTPB/SFTPA1 expression (**Figure 2g**) and accounted for 30.0% of MCseg cells versus 27.3% with 2Cseg in the same region (**Supplementary Figure S4**). In pigment-rich macrophage regions, carbon deposits reduced nuclear contrast in H&E and affected morphology-based segmentation (**Supplementary Figure S5**). Xenium measurements were also affected by optical interference of the fluorescent signals in these regions. The results illustrated complementary limitations of morphology- and fluorescence-based measurements.

The whole-transcriptome Visium HD data nonetheless allowed MCseg to identify an SPP1-positive, FTH1-positive macrophage population that could not be evaluated with a targeted panel lacking these genes. TREM2 transcripts were detected in a higher fraction of cells in this population than in the surrounding tissue (1.2%, 3 of 248 cells, versus 0.097%, 8 of 8,145 cells; **Figure 2h; Supplementary Table S6**), an observation consistent with, but based on too few positive cells to establish, the biological coherence of the assigned cell profiles.

### 3.4 External validation with expert-annotated CRC data

To assess performance beyond the LUAD development set, we next used the expert-reviewed ENACT colorectal cancer (CRC) reference containing 20,991 cell centroids, from a region of the CRC section that was not used during development. MCseg covered 65.1% of reference cells, while 34.9% fell in gaps between masks (**Figure 3a**). Coverage efficiency was 1.35, indicating that masks were preferentially concentrated over recognizable cells rather than broadly filling the tissue. Among MCseg-covered reference cells with CellTypist labels, which comprised 11,709 of the 13,673 covered cells, epithelial cells showed precision 0.89 and recall 0.92, stromal cells precision 0.78 and recall 0.59, and immune cells precision 0.38 and recall 0.54 (**Figure 3b**). The lower immune-cell performance is consistent with the small size and dense packing of lymphocytes.

**Figure 3.**
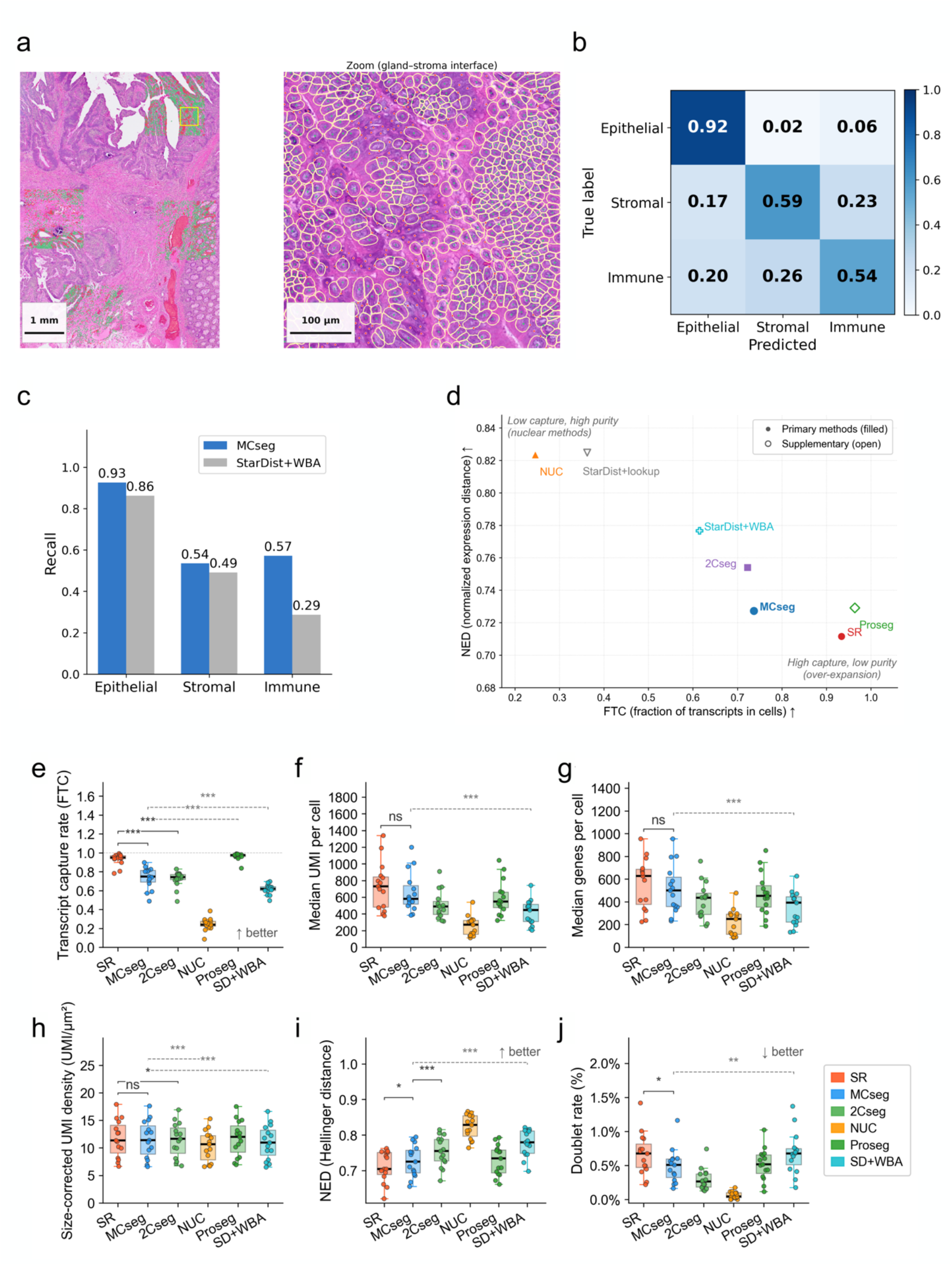
Expert-annotated CRC validation and transcript-derived benchmarking. (a) ENACT evaluation region and a representative tumor–stroma boundary. Covered reference centroids are shown in green and uncovered centroids in red. (b) Row-normalized confusion matrix for MCseg-covered cells. (c) Lineage recall for the 10,275 reference cells covered by both MCseg and the ENACT workflow. (d) FTC–NED trade-off across methods. (e–j) Boxplots across 15 CRC ROIs for FTC, median UMI count per cell, median gene count per cell, UMI density, NED, and lineage-exclusive co-expression rate. Points denote individual ROIs. Boxes show the median and interquartile range, and whiskers extend to 1.5 times the interquartile range. Significance labels in the panels are based on paired Wilcoxon signed-rank tests (ns, p > 0.05; *, p < 0.05; **, p < 0.01; ***, p < 0.001). The prespecified MCseg–SR sign-flip tests are described in the text and Table 1.

**Table 1.** Transcript assignment performance across 15 CRC ROIs.

| **Method** | **FTC** | **UMI density (UMI/µm²)** | **NED** | **Lineage-exclusive co-expression** |
|---|---|---|---|---|
| SR | 0.934 | 11.7 | 0.712 | 0.67% |
| MCseg | 0.737 | 11.6 | 0.727* | 0.49%* |
| 2Cseg | 0.723 | 11.4 | 0.754 | 0.30% |
| Proseg | 0.964 | 11.7 | 0.729 | 0.52% |
| NUC | 0.246 | 10.4 | 0.823 | 0.06% |
| StarDist | 0.362 | 10.3 | 0.825 | 0.05% |
| ENACT (StarDist + WBA) | 0.615 | 11.0 | 0.777 | 0.69% |

\* MCseg versus SR: NED p = 0.008 and lineage-exclusive co-expression p = 0.010 by sign-flip permutation test (10,000 iterations; n = 15 paired ROIs from a single tissue section).

To limit the effect of different coverage strategies, the primary direct comparison was restricted to 10,275 reference cells covered by both MCseg and the StarDist-based ENACT workflow. Within this shared set, recall was higher with MCseg for epithelial cells (0.93 versus 0.86), stromal cells (0.54 versus 0.49), and immune cells (0.57 versus 0.29). The micro-F1 scores were 0.805 and 0.723, respectively (**Figure 3c; Supplementary Table S7**). Per-class F1 under alternative coverage denominators is provided in **Supplementary Table S3**. When all 20,991 reference cells were included and uncovered cells were counted as false negatives, micro-F1 was 0.566 for MCseg and 0.708 for the ENACT workflow, and lineage-specific losses from non-coverage were 36.3% (epithelial), 37.0% (stromal), and 23.7% (immune). These results show closer agreement with the expert-reviewed lineage labels among jointly covered cells. Because the ENACT annotations themselves served as the reference standard, this comparison reflects agreement in lineage assignment rather than validation against an independent whole-cell boundary ground truth.

### 3.5 Transcript-derived benchmarking across 15 CRC regions

Because matched whole-cell reference boundaries were not available for most CRC regions, we compared seven segmentation or assignment strategies with transcript-derived measures: MCseg, SR, 2Cseg, Proseg, NUC, StarDist, and ENACT StarDist-plus-WBA workflow. SR and Proseg captured the largest fractions of tissue transcripts (FTC = 0.934 and 0.964), whereas MCseg captured 0.737 of tissue transcripts (**Table 1; per-ROI values in Supplementary Table S1**). Visual inspection in H&E images showed that SR and Proseg masks frequently extended into glandular lumina and stromal channels (**Supplementary Figure S6**). Despite the difference in FTC, UMI density was nearly identical for SR and MCseg (11.7 and 11.6 UMIs/µm^2^), and median UMI and gene counts per cell were similar (**Figure 3f, g; Supplementary Table S1**).

Relative to SR, MCseg had higher NED (0.727 versus 0.712; sign-flip permutation p = 0.008; bootstrap 95% confidence interval for the difference, 0.005-0.027) and lower lineage-exclusive co-expression (0.49% versus 0.67%; p = 0.010; bootstrap 95% confidence interval for the difference, -0.40 to -0.05 percentage points). Similar UMI density indicates comparable transcript concentration within the retained masks, but does not exclude effects of mask size, transcript capture, or cell selection on lineage-mixing estimates. These comparisons are across 15 regions of one tissue section and therefore describe within-section consistency rather than generalization across patients. Nuclear masks produced still higher NED values but captured far fewer transcripts (FTC = 0.246 for NUC and 0.362 for StarDist), and 2Cseg also showed higher NED than MCseg despite lower geometric accuracy in the LUAD. No single transcript-derived metric therefore provided a complete ranking. Instead, MCseg offers a practically balanced intermediate regime that preserved substantial transcript recovery while reducing lineage mixing relative to SR.

### 3.6 MCseg resolved lineage-distinct populations in a CRC tertiary lymphoid structure

To ask whether these quantitative differences translated into a biologically interpretable cell-level structure, we examined a tertiary lymphoid structure (TLS) candidate in the CRC section. ROI 15 was selected from three significant TIL-associated hotspot candidates identified by local Moran’s I analysis of a composite score of MS4A1, CD79A, CXCL13, IGKC, CD3E, and LTB (8,999 permutations; p < 0.01; **Supplementary Figure S8**). Within ROI 15, Leiden clustering of 636 MCseg-segmented cells resolved plasma/B-cell, stromal, myeloid, and tumor populations that formed distinct transcriptional groups and coherent spatial domains (**Figure 4; Supplementary Figure S9**). SR identified 440 cells in the same region, but its largest cluster contained 256 cells and co-expressed IGKC, VIM, and LYZ, consistent with transcript mixing across broader boundaries. JCHAIN signal was largely confined to plasma-cell-associated MCseg masks, whereas the corresponding SR masks included signal from adjacent compartments. Several gaps between MCseg masks coincided with luminal or stromal spaces visible in the H&E image rather than obvious cells. This example illustrates how segmentation choices change cell-level interpretation; it is not an independent validation of boundary accuracy.

**Figure 4.**
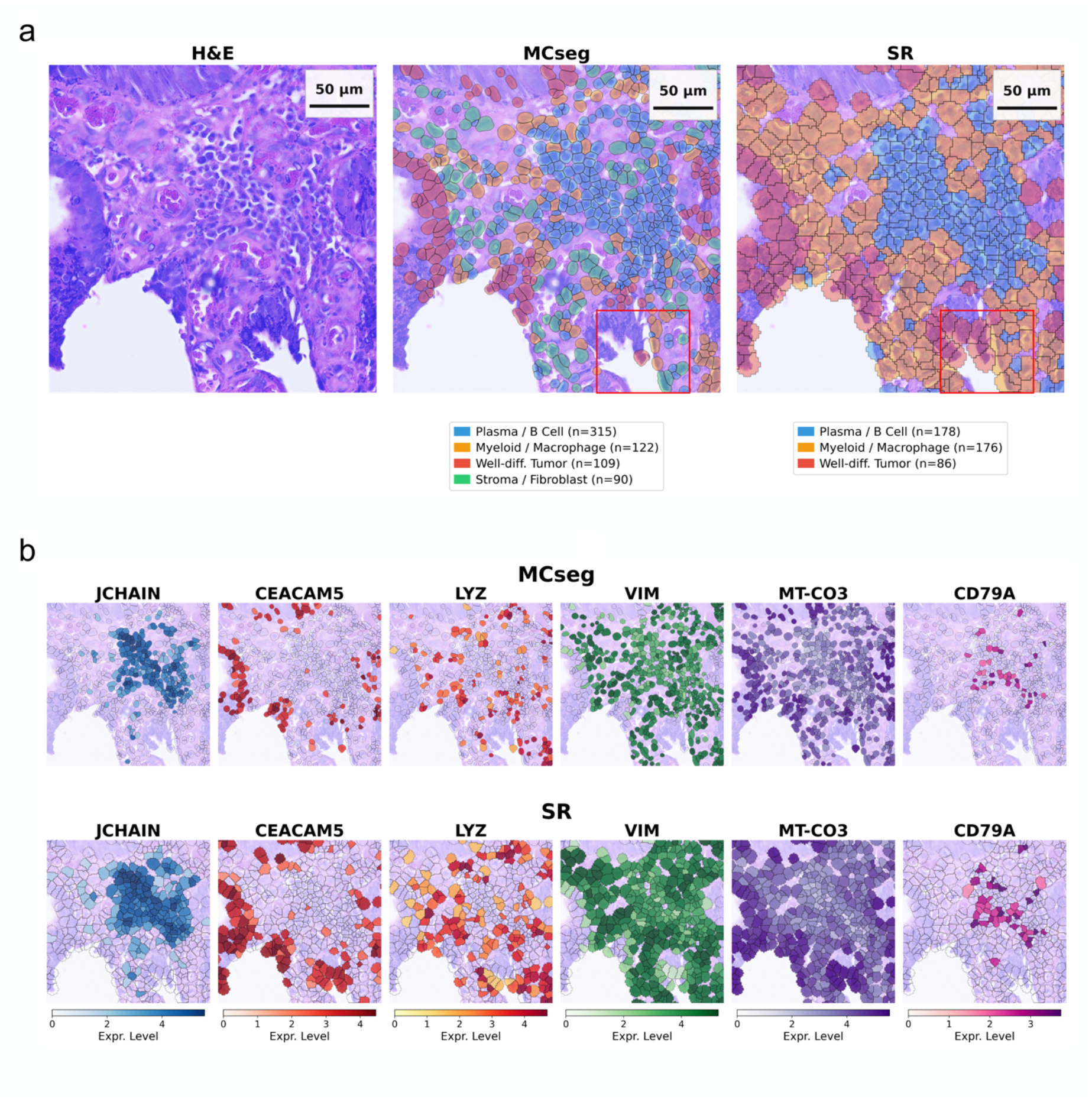
Cell-state organization in a colorectal cancer tertiary lymphoid structure. (a) H&E morphology and cell-type maps generated with MCseg and Space Ranger. (b) Spatial distribution of lineage-associated transcripts under the two segmentation strategies. MCseg produced more localized lineage signals, whereas broader Space Ranger (SR) masks showed greater overlap across neighboring compartments. Scale bars are shown in the corresponding panels.

Having established that MCseg resolved lineage-distinct populations within this structure, we then used MCseg-derived coordinates from 47,713 cells across the CRC section to test whether selected immune populations exhibited non-random spatial association. SPP1-positive macrophages (n = 391), Tregs (n = 251), and CD8-positive T cells (n = 1,509) were each closer to the other two populations than expected under complete spatial randomness. Observed versus expected median distances were 31.5 versus 45.2 µm for SPP1-positive macrophages and CD8-positive T cells, 27.0 versus 45.2 µm for Tregs and CD8-positive T cells, and 61.1 versus 90.7 µm for Tregs and SPP1-positive macrophages (all permutation p < 0.001).

A total of 53.4% of Tregs were within 30 µm of a CD8-positive T cell, compared with a median of 29% under the randomized model. Tregs in direct contact with CD8-positive T cells also had a higher CD8-positive T-cell fraction within 100 µm than non-contact Tregs (4.4% versus 3.5%; FDR = 0.002). These analyses demonstrate the type of cell-level spatial relationship that becomes accessible after transcript attribution. They describe association in a single tissue section and do not establish recruitment or migration.

### 3.7 Fixed-parameter application to breast cancer tissue

To test whether the fixed architecture can transfer beyond CRC without further tuning, we applied MCseg directly to a fresh-frozen breast cancer whole-slide dataset. MCseg segmented 96,876 cells with FTC = 0.514, median UMI count = 1,502, and NED = 0.519. The ENACT StarDist-based comparator segmented 185,483 cells with FTC = 0.612, median UMI count = 1,323, and NED = 0.560 (**Supplementary Figure S10**). These results show that the frozen MCseg workflow can be executed on a distinct tissue and sample preservation condition without a new architecture search, but they do not establish superiority in breast cancer or broad cross-tissue generalizability.

## 4. Discussion

This study shows how a limited paired spatial reference can be used to develop a segmentation and transcript-attribution workflow that is then fixed and deployed without the reference assay. The development of this platform addresses two linked challenges in cell-level spatial transcriptomics: how to design a segmentation workflow that can accommodate heterogeneous tissue morphology, and how to make that workflow accessible after development. The resulting platform separates these stages. An AI agent was used to search workflow composition against reference boundaries, whereas routine deployment uses a fixed deterministic architecture through a local no-code interface. In the LUAD development set, this fixed architecture modestly increased mean PQ over a tuned 2Cseg baseline, while the reference-guided calibration analysis showed that further agreement was possible when reference information was available for each ROI. In an expert-reviewed CRC region not used during development, MCseg showed closer agreement with lineage labels than the StarDist-based ENACT workflow among jointly covered cells. Across different CRC regions, MCseg did not maximize every metric but reduced lineage mixing relative to SR while maintaining useful transcript abundance.

### 4.1 AI-agent search as a workflow-development approach

The comparison with 2Cseg illustrates the distinction between parameter optimization and workflow search. In 2Cseg, the operation sequence was determined and fixed by the researchers, and Optuna tuned parameters within that architecture. During MCseg development, the agent could alter the composition and ordering of preprocessing, segmentation, mask integration, and expansion steps. The retained workflow therefore emerged from a broader design space than the baseline parameter search, combining complementary Cellpose models and image representations that would otherwise have required repeated manual testing. Agent-guided search identified a workflow that improved on the prespecified, conventionally tuned baseline. However, the two approaches also differed in search space, number of segmentation passes, and objective (PQ for 2Cseg versus AP@0.5 for MCseg), so this comparison does not isolate the contribution of the agent itself.

This process should not be interpreted as unconstrained autonomous discovery. The researchers selected the candidate operations, reference data, objective, prompt, execution environment, and stopping logic, and reviewed the retained code before deployment. The agent’s role was to generate, execute, score, and iteratively revise candidates within those boundaries. This is best viewed as a structured method-development strategy, not as a test of whether an AI agent is generally better than a human developer. This distinction is important because recent discussions of agentic genomics emphasize that autonomous execution shifts the bottleneck from code generation toward validation and auditability (Corpas, et al., 2026) MCseg follows that principle by exposing prompts, search records, and executable code while keeping the deployed workflow deterministic and independent of an external language model. The separation of search and deployment is important for reproducibility and lowers the barrier for users who only need the final workflow.

The same development pattern could be useful beyond this specific pipeline. Co-registered assays can provide reference-quality information during method construction while a cheaper or broader assay supplies the data available at routine deployment. Here, Xenium-derived boundaries served as a development reference for Visium HD plus H&E. Using Xenium-derived boundaries to guide an AI-agent search over segmentation and image-processing operations increased agreement with those reference boundaries without requiring Xenium data at the time of analysis. Because Xenium boundaries are themselves computational estimates and also served as the search target, this increase reflects greater agreement with Xenium-derived reference boundaries rather than attainment of an absolute level of geometric accuracy. The fixed-parameter results in Section 3.2 show a modest increase in this agreement, and the reference-guided calibration analysis indicates that further agreement is possible when sample-specific reference information is available. A more general lesson is therefore not that an AI agent replaces segmentation expertise, but that pairing an agentic AI search loop with reference-grade validation data offers a practical route to explore combinations of preprocessing, models, and post-processing to build segmentation platforms for other tissues or spatial assays that are difficult to optimize manually.

Our findings support the feasibility of using a limited paired reference to guide workflow composition rather than task-specific model training. They do not establish that agent-guided search is intrinsically more data-efficient than other optimization strategies. That question will require comparisons across reference-data budgets, matched search spaces, and independent test specimens. Likewise, demonstrating an advantage specific to the agent would require comparison with random or Bayesian search using the same operation library, objective, reference data, and a similar computational budget.

Paired and reliably registered multimodal data are costly to generate and integrate, and it is impractical to require them for every sample. The value of such data therefore need not be limited to training a new model: they can also be used to guide, evaluate, and fix an analysis workflow that other researchers can apply without the reference assay. Making this separation between development and deployment explicit, together with its demonstrated benefits and limits, is the main contribution of this study.

### 4.2 Segmentation should be evaluated with more than transcript capture

FTC is intuitive but can reward overexpansion. SR and Proseg assigned a larger fraction of tissue transcripts, yet their masks often extended into lumina and stromal spaces. Nuclear masks showed the opposite extreme: high NED and low lineage mixing but poor cytoplasmic transcript assignment. NED and lineage-exclusive co-expression therefore provided information not contained in FTC. MCseg had higher NED and lower lineage mixing than SR while maintaining nearly the same UMI density, although similar UMI density does not exclude effects of mask size, transcript capture, or cell selection on these estimates. None of these metrics is sufficient alone. NED can be inflated by conservative masks, and lineage-marker pairs can show true biological co-expression in some contexts. The higher NED values of NUC, StarDist, and 2Cseg illustrate why transcript-derived purity measures should be interpreted together with geometry, capture, and mask area rather than used as a single ranking statistic. MCseg provides a balance across these measures rather than ranking first on each one.

### 4.3 Biological and practical implications

The practical consequence of segmentation error is a distorted molecular cell profile rather than merely an imperfect outline. This was evident in the CRC TLS example, where broader SR masks produced a large cluster with mixed plasma/B-cell, stromal, and myeloid markers, whereas MCseg resolved four lineage-distinct groups with coherent spatial organization. The resulting cell coordinates then supported neighborhood analyses of Tregs, CD8-positive T cells, and SPP1-positive macrophages. The observed proximity among Tregs, CD8-positive T cells, and SPP1-positive macrophages would have been difficult to examine with 8-µm bin aggregation.

The LUAD examples also define the limits of segmentation. MCseg could recover an SPP1-positive macrophage state from whole-transcriptome Visium HD data, but better boundaries could not compensate for poor sampling of the extremely thin AT1 cytoplasm. Likewise, pigment-rich regions created optical limitations that differed between H&E and fluorescence measurements. Segmentation can improve the cellular interpretation of high-resolution spatial data, but it cannot recover molecular information that was never captured. From a deployment perspective, MCseg runs locally through a no-code interface and was evaluated on consumer-grade Apple Silicon hardware with MPS acceleration without a dedicated CUDA-enabled GPU, making ROI-scale analysis accessible to laboratories without specialized compute infrastructure.

### 4.4 Limitations and future work

There are several potential limitations of this work. The LUAD data were used for workflow development and geometric evaluation. Both the fixed-parameter and reference-guided PQ estimates are development-set results rather than independent estimates of generalization. The latter is explicitly a reference-guided calibration analysis rather than routine deployment performance, because the starting mask and expansion were selected separately for each ROI using Xenium boundaries. Co-registration between Visium HD and Xenium can also introduce small spatial offsets, and Xenium boundaries are computational estimates rather than manually drawn whole-cell contours. Additionally, the ENACT benchmark has its own bias because the reference centroids originated from StarDist before manual review. The ENACT region and the 15 CRC ROIs came from the same tissue section, although they did not overlap; statistical comparisons across the 15 ROIs therefore reflect within-section variation rather than variation across patients. In addition, the comparison with 2Cseg does not isolate the contribution of the AI agent, because search space, number of passes, and objective also differed. A stronger assessment of geometric generalizability will require fully independent datasets with matched whole-cell boundaries that were not involved in workflow development, evaluated after the workflow has been fixed. Such references need not include the full Xenium, H&E, and Visium HD combination: H&E with registered membrane staining, or manually reviewed boundaries from another section, could quantify boundary agreement, missed cells, and merges. Furthermore, transcript-derived metrics are influenced by cell size, sequencing depth, tissue composition, and true biological co-expression. They should not be interpreted without the geometric and mask-area context. Cross-tissue testing was limited to a small number of cancer specimens and did not systematically cover variation in fixation, section quality, staining, scanner characteristics, or laboratory workflow. Finally, the current workflow remains sensitive to the Voronoi expansion radius. In samples without geometric reference data, users must rely on image review and transcript-purity measures. Independent co-registered datasets, external comparison of the simpler intermediate workflow (AP@0.5 = 0.648) with the final seven-pass workflow, comparison with newly emerging segmentation models, and data-driven calibration of expansion will be important next steps. These additions will be necessary to determine how broadly the architecture generalizes beyond the tissues and acquisition settings tested here.

## Data Availability

All raw data used in this study are publicly available. The paired LUAD Xenium Prime and Visium HD dataset and the breast cancer Visium HD dataset are available from the 10x Genomics dataset portal. The CRC Visium HD dataset is available from GEO under accession GSE280318. Processed AnnData objects and segmentation masks will be deposited in Zenodo.

## Code Availability

The MCseg segmentation workflow and transcript-attribution scripts are available at https://github.com/ddmanyes/MCseg under the MIT License. The repository also contains the AutoResearch prompts, execution scripts, evaluation functions, and search history used in workflow development.

## Acknowledgements

The authors thank 10x Genomics for making the LUAD and breast cancer demonstration datasets publicly available. CRC data were obtained from GEO (GSE280318). Analyses were performed on an Apple Silicon computer with Apple MPS acceleration. The Anthropic Claude API was used during candidate workflow generation, and Claude Code assisted with code generation, figure preparation, and manuscript editing. This work was supported by National Taiwan University Hospital (NTUH114-E0008 to SJL), National Taiwan University (114L894201 to SJL), the Taiwan National Science and Technology Council (NSTC 113-2314-B-002-042-MY3 to SJL), and Academia Sinica (AS-ASSA-114-02 to SJL).

## Abbreviations

2Cseg: two-diameter Cellpose segmentation workflow
AI: artificial intelligence
AP@0.5: average precision at an intersection-over-union threshold of 0.5
AT1: alveolar type 1
AT2: alveolar type 2
CLAHE: contrast-limited adaptive histogram equalization
CRC: colorectal cancer
CUDA: Compute Unified Device Architecture
DCIS: ductal carcinoma in situ
ENACT: end-to-end analysis of Visium High Definition data
FDR: false discovery rate
FFPE: formalin-fixed, paraffin-embedded
FTC: fractional transcript capture
GEO: Gene Expression Omnibus
GPU: graphics processing unit
H&E: hematoxylin and eosin
HD: high definition
HED: hematoxylin–eosin–DAB
HVG: highly variable gene
LUAD: lung adenocarcinoma
MCseg: Multiple Cellpose Segmentation
MPS: Metal Performance Shaders
MWU: Mann–Whitney U
NED: neighborhood expression discordance
NUC: nucleus-restricted segmentation
PCA: principal component analysis
PQ: panoptic quality
RQ: recognition quality
ROI: region of interest
SQ: segmentation quality
ST: spatial transcriptomics
TLS: tertiary lymphoid structure
TPE: tree-structured Parzen estimator
Treg: regulatory T cell
UMI: unique molecular identifier
UMAP: Uniform Manifold Approximation and Projection
WBA: weighted bin-area assignment

