## Supplemental Data for "MCseg: reference-guided AI-agent development of a fixed cell segmentation and transcript attribution workflow for spatial transcriptomics"

**Supplementary materials**

### Supplementary Note 1: MCseg algorithm specification and AI-agent development environment

#### A. AI-agent development environment

MCseg was developed through iterative AI-agent–guided optimization within a researcher-defined sandbox. The agent modified a single executable segmentation script and evaluated candidate workflows against Xenium-derived reference masks using average precision at an intersection-over-union threshold of 0.5 (AP@0.5). The researchers defined the candidate tool library, reference data, scoring function, prompt, execution limit, and retained-code policy, and reviewed the converged configuration before deployment. This arrangement separated automated candidate generation from the scientific decisions that defined the search space and final workflow.

The candidate library included Cellpose cyto2, cyto3, nuclei, and cpsam models; contrast-limited adaptive histogram equalization (CLAHE); hematoxylin extraction by Ruifrok–Johnston H&E color deconvolution; Gaussian smoothing; watershed; Voronoi tessellation; and morphological operations. The search space covered model selection and combination, diameter settings, CLAHE parameters, cell-probability and flow thresholds, ensemble overlap thresholds, expansion strategy, and area filters.

The optimization log contained both successful and failed candidate executions; therefore, no single cycle count was used as a scientific endpoint. On the primary development patch, AP@0.5 increased from 0.320 for the initial single-model workflow to 0.650 for the retained architecture. This score describes the development patch and is distinct from the six-ROI fixed-parameter deployment benchmark reported in the main text. Most of this gain accompanied the introduction of CLAHE preprocessing and Voronoi-constrained expansion (AP@0.5 = 0.648); the retained seven-pass ensemble added a further 0.002 on this patch (Supplementary Figure S11).

The AI agent was used only during development. Routine MCseg analysis uses the consolidated deterministic workflow and does not call an external language model. The geometric analyses reported in the study were accelerated with Apple MPS, while the software also supports CPU-only execution. CPU runtime was not benchmarked systematically. The following specification therefore describes the fixed deployment engine rather than the development-time search loop.

#### B. MCseg pipeline specification

The fixed platform processes an H&E image together with spatial expression data and produces a labelled cell mask and a cell-by-gene expression matrix. The seven steps below describe the order used in deployment.

##### Step 1 — Image preprocessing

- CLAHE contrast enhancement (clip limit 3.0; tile grid 8 × 8).
- Hematoxylin-channel extraction by Ruifrok–Johnston H&E color deconvolution, followed by CLAHE.
- Generation of a tissue mask to prevent expansion into background regions.

**Step 2 — Seven-pass Cellpose detection**

| Pass | Model | Input | Diameter (px) | Flow threshold | Cell-probability threshold | Augment |
|---|---|---|---|---|---|---|
| 1 | cyto3 | CLAHE RGB | 17 | 0.4 | −2.0 | Yes |
| 2 | cyto3 | Hematoxylin | 17 | 0.4 | −2.0 | Yes |
| 3 | cyto3 | CLAHE RGB | 22 | 0.5 | −1.0 | No |
| 4 | cyto3 | CLAHE RGB | 13 | 0.4 | −3.0 | No |
| 5 | cpsam | CLAHE RGB | 30 | 0.4 | −1.0 | No |
| 6 | cpsam | CLAHE RGB | 16 | 0.4 | −2.0 | No |
| 7 | cpsam | Hematoxylin | 30 | 0.4 | −1.0 | No |

**Step 3 — Priority-based ensemble merging**

In deployment mode, the cyto3 17-pixel CLAHE-RGB mask initializes the ensemble. Masks from the remaining passes are considered sequentially. A candidate object is added only when less than 15% of its pixels overlap the occupied ensemble mask; only its previously unoccupied pixels are retained. This rule preserves complementary detections while limiting redundant proposals. In the reference-guided calibration analysis, the best-performing single-pass starting mask was selected using the reference mask before the same merging procedure was applied.

**Step 4 — Optional transcript-density rescue**

When spatial expression coordinates are supplied, smoothed transcript-density peaks that are not covered by an existing mask can be used to seed additional small masks. This optional step was not used in the H&E-only geometric benchmark in which the transcript input was set to null.

**Step 5 — Voronoi-constrained boundary expansion**

Unassigned tissue pixels are allocated to the nearest detected cell up to a fixed, profile-defined maximum distance, preventing overlap between adjacent labels. The deployed platform uses fixed defaults that can be adjusted by the user or by tissue profile. For the separate reference-guided calibration analysis, expansion method and distance were selected independently for each ROI using the Xenium masks; these calibration results are reported separately from fixed-parameter deployment and are not a strict upper bound.

**Step 6 — Quality filtering**

Masks are filtered by area after merging and expansion. The implementation removes small fragments and very large objects using stage-specific area thresholds; the final geometric benchmark used a retained area range of 20–6000 pixels$^2$.

**Step 7 — Transcript attribution**

Each in-tissue 2-µm Visium HD bin is assigned to the segmentation mask containing its full-resolution pixel centroid. Full-slide bin coordinates are transformed to ROI-local coordinates before mask lookup. Bins whose centroids fall outside all masks receive label 0 and are treated as unassigned. Gene counts from bins sharing the same positive mask label are summed by sparse matrix multiplication to generate the cell-by-gene AnnData matrix. This centroid-lookup rule is distinct from ENACT weighted bin-area assignment.

### Supplementary Note 2: Segmentation strategy selection and comparator models

To place MCseg in context, three supplementary comparators—Proseg, NUC, and StarDist with ENACT (A–C)—were evaluated in addition to Space Ranger and 2Cseg. Development of the 2Cseg baseline itself is described in D.

#### A. Proseg

Proseg (Jones, et al., 2025) was designed primarily for point-based spatial data. For the CRC comparison, the 2-µm bin matrix was expanded into pseudo-transcript coordinates by placing each count at the bin centroid with uniform sub-bin jitter, and the resulting file was processed with Proseg v3.1.0 across 15 ROIs. Proseg was used as a supplementary RNA-assignment comparator rather than a matched geometric benchmark.

#### B. NUC

NUC served as a conservative nuclear lower-bound control and used a single Cellpose nuclei-model pass without CLAHE or Voronoi expansion.

#### C. StarDist and ENACT

StarDist nuclear masks with centroid lookup were included as a conservative image-based comparator. The ENACT workflow combined StarDist-derived masks with neighboring weighted bin-area assignment (WBA) and was evaluated separately (Kamel, et al., 2025).

#### D. Development of the 2Cseg baseline

2Cseg was defined as a dual-diameter Cellpose nuclei workflow with CLAHE preprocessing and post-segmentation expansion. Parameters were optimized with Optuna Bayesian optimization (TPE sampler; 50 trials) on three LUAD ROIs using mean PQ against Xenium boundaries as the objective. The selected configuration used dia_small = 9.93 µm, dia_large = 35.77 µm, and 6-pixel dilation, together with the optimized flow, cell-probability, fragment, and contrast parameters. Evaluation across six LUAD ROIs yielded mean PQ = 0.432 ± 0.037. The Optuna analysis defines performance within the researcher-specified 2Cseg architecture and does not constitute an exhaustive search over alternative workflow compositions.

### Supplementary Note 3: Spatial transcriptomic datasets

#### Lung adenocarcinoma (LUAD)

The datasets served different roles in the study: LUAD supported workflow development and geometric analysis, CRC supported external and transcript-derived evaluation, and breast cancer provided a fixed-architecture transfer test. For LUAD, data were obtained from a publicly available 10x Genomics demonstration dataset comprising co-registered Visium HD (2 μm/bin) and Xenium Prime measurements from the same FFPE lung adenocarcinoma section. Six fixed-size ROIs spanning tumor and alveolar microenvironments were used for geometric analysis. Additional macrophage-rich and normal-alveolar regions were used for biological illustration. Xenium cell boundaries from cell_boundaries.parquet served as the geometric reference. Because these registered data were used during workflow development, the geometric analyses are development-set evaluations.

#### Colorectal cancer (CRC)

The CRC Visium HD FFPE dataset was obtained from GEO (GSE280318). Fifteen ROIs were selected across tumor, stroma, glandular, and immune-rich regions. Because all 15 ROIs come from one tissue section, they are within-section replicates rather than independent biological specimens. ROI 15 was selected as a tertiary lymphoid structure candidate after combining a composite score based on MS4A1, CD79A, CXCL13, IGKC, CD3E, and LTB with local Moran's I spatial autocorrelation. The ENACT expert-reviewed reference comprised 20,991 cell centroids from a non-overlapping region of the same public section [7].

#### Breast cancer

The breast cancer dataset was the publicly available 10x Genomics 'Visium HD Spatial Gene Expression Library, Human Breast Cancer (Fresh Frozen)' demonstration dataset containing ductal carcinoma in situ. It was not an ENACT dataset. MCseg was applied to the full in-tissue crop without breast-specific architecture search, using the fixed profile reported in Supplementary Figure S10. This analysis was intended as a transfer test rather than a comparative claim of superiority.

### Supplementary Tables

**Supplementary Table S1.** Per-ROI transcript attribution metrics across 15 CRC ROIs and six segmentation methods; per-ROI values for the ENACT workflow (StarDist + WBA) reported in main-text Table are not included. FTC, fraction of in-tissue UMIs assigned to cell masks; UMI density, assigned UMIs per total mask area (UMI/$\mu m^2$); NED, neighborhood expression discordance (Hellinger distance, range 0–1; higher values indicate greater separation between adjacent cell profiles); doublet rate, lineage-exclusive co-expression rate averaged across four marker pairs; N cells, segmented cells per ROI. Proseg and StarDist use centroid lookup attribution in this table.

| Method | ROI | FTC | UMI density | Median UMI | Median genes | NED | Doublet rate | N cells |
|---|---|---|---|---|---|---|---|---|
| Space Ranger | ROI 1 | 0.992 | 8.16 | 430 | 322 | 0.750 | 0.0045 | 1281 |
| MCseg | ROI 1 | 0.853 | 8.18 | 465 | 354 | 0.745 | 0.0061 | 1032 |
| 2Cseg | ROI 1 | 0.780 | 8.09 | 350 | 272 | 0.759 | 0.0027 | 1219 |
| NUC | ROI 1 | 0.208 | 6.86 | 113 | 92 | 0.840 | 0.0000 | 836 |
| Proseg | ROI 1 | 0.958 | 9.27 | 391 | 308 | 0.757 | 0.0044 | 1318 |
| StarDist+lookup | ROI 1 | 0.358 | 9.04 | 142 | 113 | 0.824 | 0.0000 | 1244 |
| Space Ranger | ROI 2 | 0.971 | 10.19 | 476 | 339 | 0.704 | 0.0026 | 1260 |
| MCseg | ROI 2 | 0.826 | 10.05 | 500 | 366 | 0.701 | 0.0028 | 1054 |
| 2Cseg | ROI 2 | 0.778 | 10.05 | 407 | 301 | 0.726 | 0.0014 | 1241 |
| NUC | ROI 2 | 0.239 | 9.47 | 166 | 130 | 0.814 | 0.0003 | 820 |
| Proseg | ROI 2 | 0.960 | 11.52 | 472 | 338 | 0.715 | 0.0032 | 1253 |
| StarDist+lookup | ROI 2 | 0.364 | 11.22 | 183 | 139 | 0.775 | 0.0002 | 1176 |
| Space Ranger | ROI 3 | 0.935 | 7.30 | 399 | 226 | 0.653 | 0.0041 | 1098 |
| MCseg | ROI 3 | 0.814 | 6.89 | 394 | 232 | 0.655 | 0.0022 | 1008 |
| 2Cseg | ROI 3 | 0.712 | 7.13 | 313 | 188 | 0.672 | 0.0019 | 1160 |
| NUC | ROI 3 | 0.216 | 7.05 | 138 | 85 | 0.764 | 0.0010 | 737 |
| Proseg | ROI 3 | 0.969 | 8.11 | 342 | 194 | 0.658 | 0.0035 | 1286 |
| StarDist+lookup | ROI 3 | 0.345 | 9.17 | 167 | 97 | 0.701 | 0.0013 | 970 |
| Space Ranger | ROI 4 | 0.960 | 15.52 | 796 | 665 | 0.741 | 0.0101 | 1067 |
| MCseg | ROI 4 | 0.812 | 15.60 | 970 | 801 | 0.725 | 0.0116 | 795 |
| 2Cseg | ROI 4 | 0.771 | 15.20 | 723 | 611 | 0.755 | 0.0075 | 973 |
| NUC | ROI 4 | 0.275 | 13.78 | 334 | 299 | 0.829 | 0.0012 | 648 |
| Proseg | ROI 4 | 0.983 | 16.30 | 831 | 682 | 0.751 | 0.0103 | 1023 |

| Method | ROI | FTC | UMI density | Median UMI | Median genes | NED | Doublet rate | N cells |
|---|---|---|---|---|---|---|---|---|
| StarDist+lookup | ROI 4 | 0.376 | 15.31 | 324 | 294 | 0.851 | 0.0010 | 984 |
| Space Ranger | ROI 5 | 0.973 | 14.77 | 857 | 680 | 0.700 | 0.0058 | 1043 |
| MCseg | ROI 5 | 0.804 | 14.75 | 1010 | 796 | 0.698 | 0.0060 | 792 |
| 2Cseg | ROI 5 | 0.744 | 14.13 | 715 | 582 | 0.736 | 0.0028 | 993 |
| NUC | ROI 5 | 0.276 | 12.22 | 381 | 334 | 0.806 | 0.0000 | 607 |
| Proseg | ROI 5 | 0.990 | 16.16 | 938 | 748 | 0.711 | 0.0044 | 968 |
| StarDist+lookup | ROI 5 | 0.354 | 14.13 | 298 | 265 | 0.829 | 0.0002 | 1001 |
| Space Ranger | ROI 6 | 0.943 | 12.75 | 732 | 627 | 0.752 | 0.0022 | 1017 |
| MCseg | ROI 6 | 0.898 | 12.74 | 602 | 522 | 0.781 | 0.0017 | 1204 |
| 2Cseg | ROI 6 | 0.830 | 12.47 | 500 | 438 | 0.806 | 0.0013 | 1373 |
| NUC | ROI 6 | 0.387 | 11.10 | 306 | 281 | 0.867 | 0.0003 | 983 |
| Proseg | ROI 6 | 0.968 | 13.12 | 492 | 432 | 0.808 | 0.0012 | 1487 |
| StarDist+lookup | ROI 6 | 0.448 | 11.46 | 318 | 292 | 0.858 | 0.0002 | 1116 |
| Space Ranger | ROI 7 | 0.958 | 17.94 | 962 | 785 | 0.704 | 0.0050 | 1008 |
| MCseg | ROI 7 | 0.802 | 17.62 | 701 | 583 | 0.751 | 0.0038 | 1173 |
| 2Cseg | ROI 7 | 0.771 | 16.93 | 524 | 455 | 0.788 | 0.0018 | 1556 |
| NUC | ROI 7 | 0.235 | 15.30 | 271 | 250 | 0.860 | 0.0012 | 841 |
| Proseg | ROI 7 | 0.976 | 18.50 | 606 | 518 | 0.771 | 0.0020 | 1515 |
| StarDist+lookup | ROI 7 | 0.392 | 17.13 | 284 | 258 | 0.855 | 0.0007 | 1386 |
| Space Ranger | ROI 8 | 0.982 | 7.86 | 491 | 415 | 0.762 | 0.0068 | 1000 |
| MCseg | ROI 8 | 0.675 | 7.75 | 556 | 460 | 0.763 | 0.0056 | 666 |
| 2Cseg | ROI 8 | 0.731 | 7.48 | 430 | 364 | 0.787 | 0.0039 | 893 |
| NUC | ROI 8 | 0.238 | 6.70 | 198 | 182 | 0.854 | 0.0004 | 560 |

| Method | ROI | FTC | UMI density | Median UMI | Median genes | NED | Doublet rate | N cells |
|---|---|---|---|---|---|---|---|---|
| Proseg | ROI 8 | 0.981 | 8.56 | 523 | 434 | 0.773 | 0.0067 | 971 |
| StarDist+lookup | ROI 8 | 0.381 | 7.88 | 182 | 167 | 0.860 | 0.0003 | 945 |
| Space Ranger | ROI 9 | 0.945 | 12.68 | 773 | 651 | 0.753 | 0.0074 | 945 |
| MCseg | ROI 9 | 0.749 | 12.83 | 635 | 555 | 0.794 | 0.0043 | 983 |
| 2Cseg | ROI 9 | 0.755 | 12.53 | 489 | 436 | 0.816 | 0.0022 | 1228 |
| NUC | ROI 9 | 0.295 | 12.27 | 328 | 298 | 0.863 | 0.0003 | 770 |
| Proseg | ROI 9 | 0.987 | 12.73 | 550 | 480 | 0.802 | 0.0051 | 1286 |
| StarDist+lookup | ROI 9 | 0.388 | 12.11 | 286 | 261 | 0.866 | 0.0002 | 1067 |
| Space Ranger | ROI 10 | 0.897 | 6.64 | 377 | 238 | 0.684 | 0.0089 | 894 |
| MCseg | ROI 10 | 0.707 | 6.60 | 385 | 246 | 0.686 | 0.0059 | 805 |
| 2Cseg | ROI 10 | 0.676 | 6.73 | 319 | 207 | 0.702 | 0.0038 | 930 |
| NUC | ROI 10 | 0.233 | 6.59 | 133 | 91 | 0.780 | 0.0004 | 625 |
| Proseg | ROI 10 | 0.978 | 8.38 | 381 | 242 | 0.694 | 0.0078 | 998 |
| StarDist+lookup | ROI 10 | 0.317 | 8.75 | 137 | 96 | 0.755 | 0.0006 | 847 |
| Space Ranger | ROI 11 | 0.950 | 9.95 | 670 | 418 | 0.661 | 0.0091 | 854 |
| MCseg | ROI 11 | 0.572 | 9.62 | 535 | 328 | 0.668 | 0.0051 | 690 |
| 2Cseg | ROI 11 | 0.723 | 9.98 | 449 | 312 | 0.707 | 0.0038 | 1054 |
| NUC | ROI 11 | 0.185 | 8.61 | 168 | 118 | 0.782 | 0.0009 | 586 |
| Proseg | ROI 11 | 0.981 | 11.72 | 528 | 376 | 0.703 | 0.0052 | 1156 |
| StarDist+lookup | ROI 11 | 0.355 | 11.52 | 206 | 144 | 0.774 | 0.0010 | 1006 |
| Space Ranger | ROI 12 | 0.783 | 13.49 | 696 | 590 | 0.755 | 0.0050 | 800 |
| MCseg | ROI 12 | 0.589 | 13.30 | 579 | 499 | 0.787 | 0.0037 | 815 |
| 2Cseg | ROI 12 | 0.586 | 13.08 | 504 | 445 | 0.802 | 0.0016 | 926 |

| Method | ROI | FTC | UMI density | Median UMI | Median genes | NED | Doublet rate | N cells |
|---|---|---|---|---|---|---|---|---|
| NUC | ROI 12 | 0.252 | 11.82 | 314 | 284 | 0.855 | 0.0008 | 650 |
| Proseg | ROI 12 | 0.838 | 12.14 | 562 | 485 | 0.786 | 0.0065 | 959 |
| StarDist+lookup | ROI 12 | 0.304 | 12.22 | 284 | 260 | 0.858 | 0.0003 | 836 |
| Space Ranger | ROI 13 | 0.971 | 15.39 | 1191 | 954 | 0.695 | 0.0071 | 773 |
| MCseg | ROI 13 | 0.744 | 15.10 | 1199 | 955 | 0.711 | 0.0053 | 655 |
| 2Cseg | ROI 13 | 0.770 | 14.52 | 911 | 759 | 0.744 | 0.0029 | 849 |
| NUC | ROI 13 | 0.297 | 13.31 | 540 | 479 | 0.789 | 0.0010 | 521 |
| Proseg | ROI 13 | 0.987 | 15.88 | 1052 | 858 | 0.730 | 0.0061 | 865 |
| StarDist+lookup | ROI 13 | 0.371 | 14.42 | 449 | 407 | 0.836 | 0.0009 | 797 |
| Space Ranger | ROI 14 | 0.939 | 11.14 | 835 | 694 | 0.736 | 0.0074 | 776 |
| MCseg | ROI 14 | 0.722 | 11.40 | 780 | 652 | 0.753 | 0.0073 | 684 |
| 2Cseg | ROI 14 | 0.724 | 11.22 | 579 | 499 | 0.780 | 0.0046 | 867 |
| NUC | ROI 14 | 0.265 | 9.75 | 316 | 284 | 0.838 | 0.0018 | 567 |
| Proseg | ROI 14 | 0.961 | 12.25 | 676 | 571 | 0.775 | 0.0068 | 914 |
| StarDist+lookup | ROI 14 | 0.365 | 10.94 | 270 | 244 | 0.865 | 0.0011 | 887 |
| Space Ranger | ROI 15 | 0.807 | 11.35 | 1340 | 821 | 0.621 | 0.0142 | 440 |
| MCseg | ROI 15 | 0.488 | 11.07 | 547 | 376 | 0.690 | 0.0024 | 636 |
| 2Cseg | ROI 15 | 0.487 | 11.63 | 381 | 280 | 0.729 | 0.0023 | 885 |
| NUC | ROI 15 | 0.085 | 10.64 | 145 | 114 | 0.813 | 0.0000 | 327 |
| Proseg | ROI 15 | 0.941 | 13.09 | 664 | 454 | 0.686 | 0.0058 | 855 |
| StarDist+lookup | ROI 15 | 0.311 | 13.86 | 205 | 160 | 0.789 | 0.0005 | 932 |

**Supplementary Table S2.** Selected LUAD parent regions used to define benchmark or biological-validation areas. Coordinates refer to broad tissue regions in the full-resolution H&E image (0.2737 µm/pixel), not to the six fixed-size geometric benchmark crops.

| Region | Description | x (px) | y (px) | Width (px) | Height (px) | Width (µm) | Height (µm) |
|---|---|---|---|---|---|---|---|
| Tumor boundary | G2/G3 grade transition zone with lymphoid infiltration | 9202 | 16552 | 6022 | 4521 | 1648 | 1238 |
| HEV / TLS | High endothelial venule cluster within tertiary lymphoid structure | 11787 | 17937 | 1402 | 1151 | 384 | 315 |
| Dust cells | Carbon-laden alveolar macrophage aggregate | 7934 | 11264 | 2869 | 1602 | 785 | 438 |
| Normal lung | Open alveolar region; AT1/AT2 pneumocyte validation | 7050 | 23275 | 2069 | 1267 | 566 | 347 |

**Supplementary Table S3.** Per-class F1 among ENACT reference cells covered by MCseg. These subset analyses use MCseg-specific covered-cell denominators (11,709 cells for lookup and 11,948 for WBA) and are distinct from the jointly covered comparison in the main text (10,275 cells) and from full-denominator analyses in which uncovered reference cells are false negatives.

| Cell type | MCseg + lookup | MCseg + WBA |
|---|---|---|
| Epithelial | 0.91 | 0.91 |
| Stromal | 0.68 | 0.68 |
| Immune | 0.45 | 0.49 |
| **Micro F1 (all classes)** | **0.790** | **0.802** |

**Supplementary Table S4.** Fixed-parameter deployment benchmark: MCseg versus 2Cseg across six LUAD ROIs. The MCseg masks were generated without access to Xenium boundaries; reference masks were used only for scoring. Standard deviation is calculated across ROIs (population SD, matching the analysis output).

| ROI | Tissue | MCseg PQ | MCseg SQ | MCseg RQ | 2Cseg PQ | 2Cseg SQ | 2Cseg RQ |
|---|---|---|---|---|---|---|---|
| ROI 1 | Tumor boundary | 0.530 | 0.765 | 0.693 | 0.502 | 0.643 | 0.782 |
| ROI 2 | Tumor stroma | 0.422 | 0.711 | 0.594 | 0.438 | 0.596 | 0.734 |
| ROI 3 | Mixed | 0.508 | 0.754 | 0.673 | 0.401 | 0.550 | 0.729 |
| ROI 4 | Normal–tumor interface | 0.437 | 0.719 | 0.608 | 0.409 | 0.556 | 0.736 |
| ROI 5 | Alveolar | 0.360 | 0.680 | 0.530 | 0.395 | 0.551 | 0.716 |
| ROI 6 | Tumor core | 0.572 | 0.786 | 0.728 | 0.450 | 0.614 | 0.733 |
| **Mean ± SD** | | **0.472 ± 0.072** | **0.736** | **0.638** | **0.432 ± 0.037** | **0.585** | **0.739** |

**Supplementary Table S5.** Reference-guided calibration analysis across six LUAD ROIs. For this analysis, the single-pass starting mask, expansion strategy, and expansion distance were selected independently for each ROI using the Xenium mask. These values describe sample-specific calibration with access to the reference; they are not a strict upper bound and are not routine deployment results.

| ROI | Tissue | MCseg PQ | MCseg SQ | MCseg RQ | 2Cseg PQ | 2Cseg SQ | 2Cseg RQ |
|---|---|---|---|---|---|---|---|
| ROI 1 | Tumor boundary | 0.660 | 0.830 | 0.795 | 0.502 | 0.643 | 0.782 |
| ROI 2 | Tumor stroma | 0.549 | 0.775 | 0.709 | 0.438 | 0.596 | 0.734 |
| ROI 3 | Mixed | 0.523 | 0.762 | 0.687 | 0.401 | 0.550 | 0.729 |
| ROI 4 | Normal–tumor interface | 0.470 | 0.735 | 0.639 | 0.409 | 0.556 | 0.736 |
| ROI 5 | Alveolar | 0.514 | 0.757 | 0.679 | 0.395 | 0.551 | 0.716 |
| ROI 6 | Tumor core | 0.608 | 0.804 | 0.756 | 0.450 | 0.614 | 0.733 |
| **Mean ± SD** | | **0.554 ± 0.063** | **0.777** | **0.711** | **0.432 ± 0.037** | **0.585** | **0.739** |

**Supplementary Table S6.** TREM2 transcript detection in the SPP1$^+$/FTH1$^+$ macrophage cluster (C4) versus background, ROI9 (LUAD macrophage-rich region). "% positive" denotes the binary detection rate — the percentage of cells assigned at least one TREM2 transcript — and is independent of per-cell expression level. TREM2-positive cell counts were derived from the ROI9 cell-by-gene matrix; the all-cell positivity rate (0.13%) matches the recorded CSV summary value.

| Cell Population | Cells (n) | TREM2-positive cells (n) | % positive |
|---|---|---|---|
| C4 (SPP1$^+$/FTH1$^+$ macrophage cluster) | 248 | 3 | 1.2% |
| Remaining ROI9 cells (background) | 8,145 | 8 | 0.097% |
| All ROI9 cells | 8,393 | 11 | 0.13% |

**Supplementary Table S7.** Per-class precision, recall, and F1 for MCseg versus the StarDist-based ENACT workflow (StarDist+WBA), restricted to the 10,275 ENACT reference cells jointly covered by both methods (main text Section 3.4). Micro F1 (all classes) is computed across all three cell types combined.

| Cells type | Support (n) | MCseg precision | MCseg recall | MCseg F1 | StarDist+WBA precision | StarDist+WBA recall | StarDist+WBA F1 |
|---|---|---|---|---|---|---|---|
| Epithelial | 6,993 | 0.903 | 0.926 | 0.915 | 0.809 | 0.863 | 0.835 |
| Stromal | 2,192 | 0.779 | 0.535 | 0.635 | 0.747 | 0.492 | 0.593 |
| Immune | 1,090 | 0.390 | 0.572 | 0.464 | 0.228 | 0.287 | 0.254 |
| **Macro F1 (all classes)** | **10,275** | **0.805** | | | **0.723** | | |

### Supplementary Figures

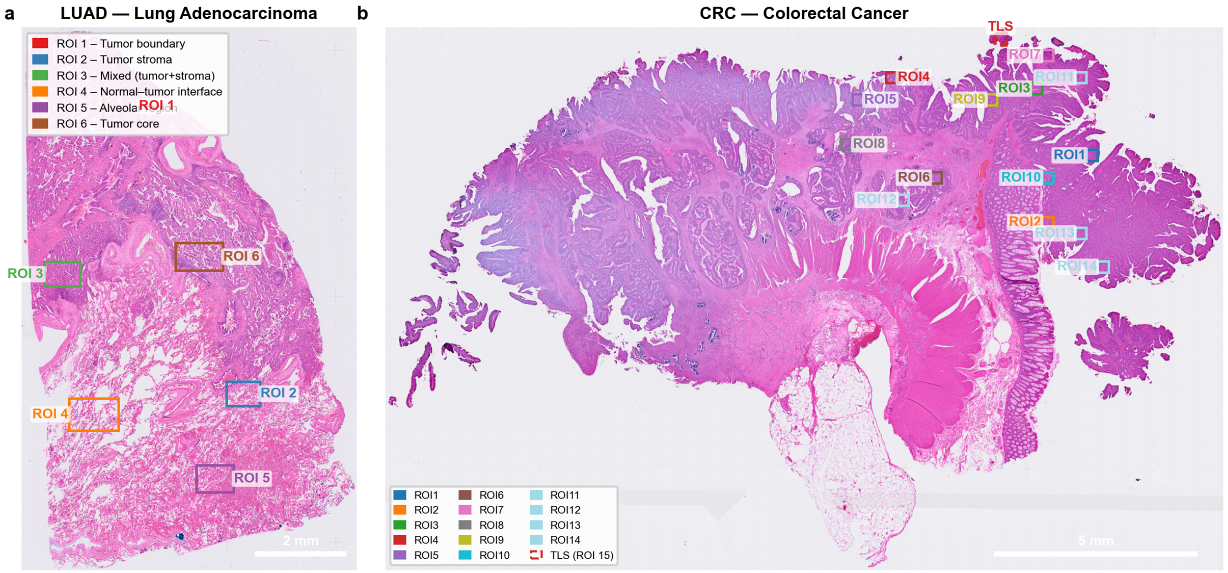


**Supplementary Figure S1.** ROI location overview on full-slide H&E. (a) LUAD section showing the six ROIs used for geometric benchmarking. (b) CRC section showing the 15 ROIs used for transcript-derived benchmarking; ROI 15, selected for TLS analysis, is highlighted by the dashed red box. Scale bars: 2 mm (LUAD) and 5 mm (CRC).

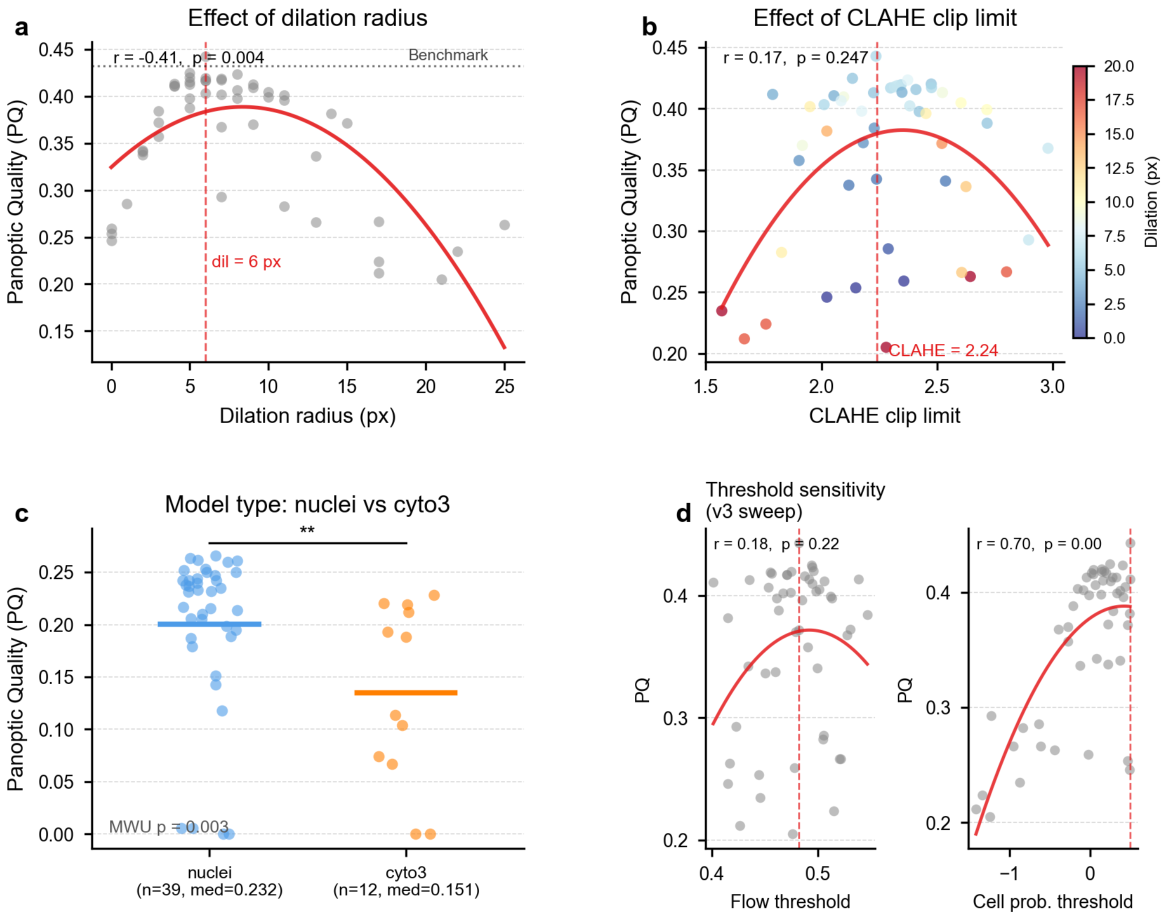


**Supplementary Figure S2.** 2Cseg hyperparameter sensitivity analysis. Sensitivity of PQ to selected parameters from the 50-trial Optuna search across three LUAD ROIs. (a) Dilation radius. (b) CLAHE clip limit, with dilation radius indicated by color. (c) Cellpose model type. (d) Flow and cell-probability thresholds. The results indicated limited additional improvement within the predefined 2Cseg parameter space and motivated exploration of alternative workflow compositions. MWU, Mann–Whitney U test; values below the displayed precision are reported as $p < 0.001$.

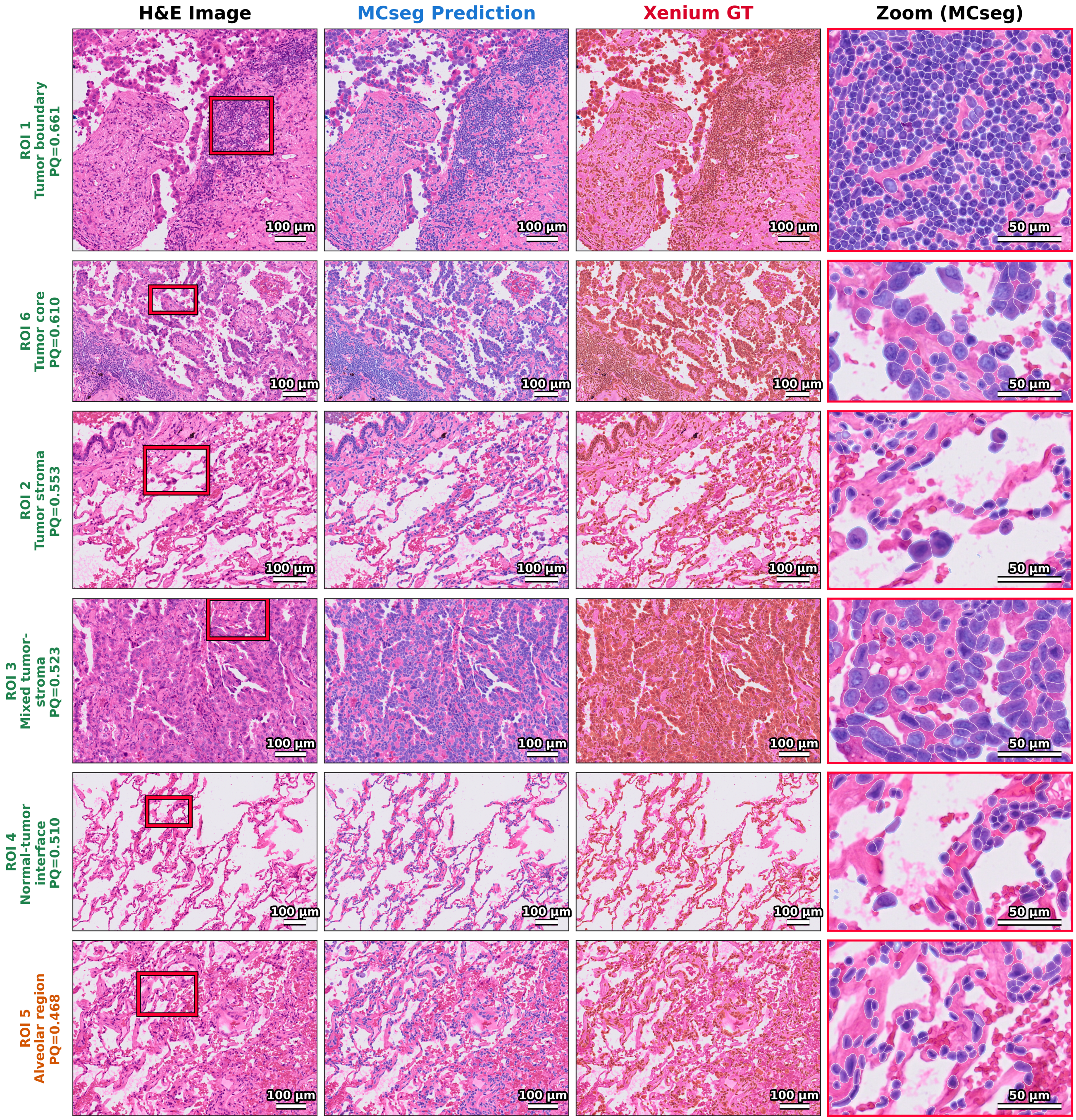


**Supplementary Figure S3.** Reference-guided MCseg upper-bound masks and Xenium reference boundaries across six LUAD ROIs. Each row shows the H&E image, the MCseg mask generated with ROI-specific reference-guided expansion selection, the Xenium reference mask, and a representative zoomed region. PQ is annotated for each ROI. These panels illustrate the upper-bound analysis in Supplementary Table S5 and should not be interpreted as fixed-parameter deployment. Scale bars: 50 µm.

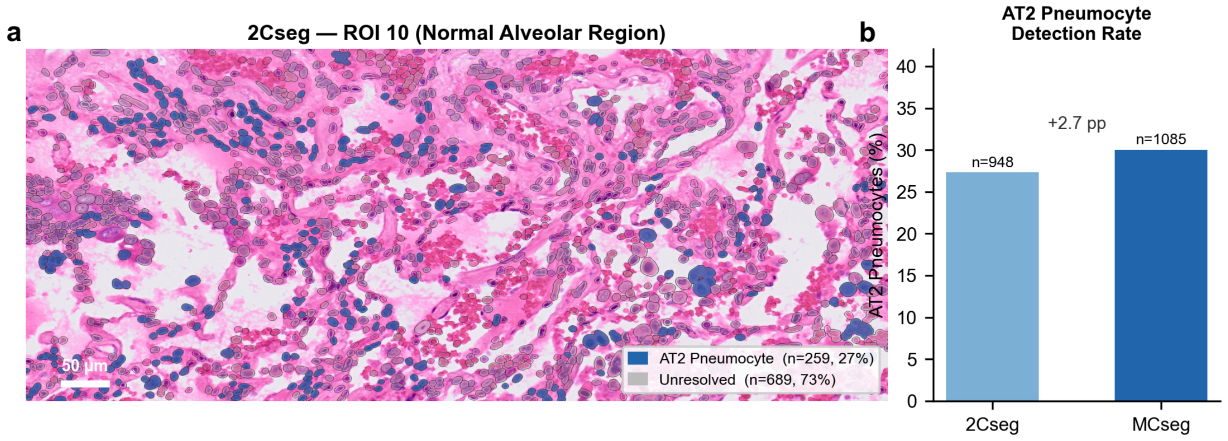


**Supplementary Figure S4.** LUAD normal-alveolar region: AT2-associated cell detection with 2Cseg and MCseg. (a) Leiden-based 2Cseg cell map, with the AT2-associated cluster highlighted in blue (259 of 948 cells; 27.3%). (b) The corresponding AT2-associated cluster accounted for 30.0% of MCseg cells (326 of 1,085), an increase of 2.7 percentage points. These percentages describe cluster assignment rather than a direct measurement of cell-recovery sensitivity. Scale bar: 50 µm.

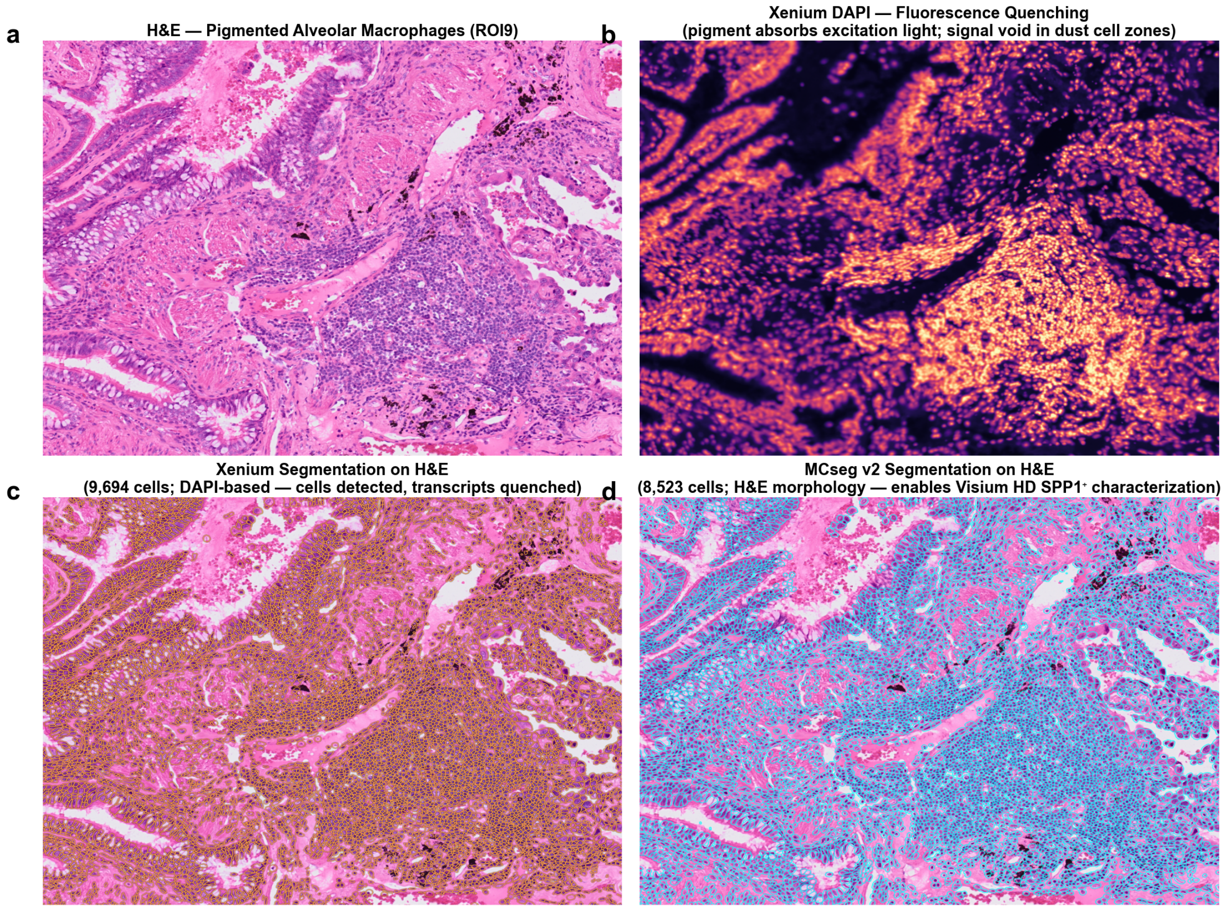


**Supplementary Figure S5.** LUAD normal-alveolar region: AT2-associated cell detection with 2Cseg and MCseg. (a) Leiden-based 2Cseg cell map, with the AT2-associated cluster highlighted in blue (259 of 948 cells; 27.3%). (b) The corresponding AT2-associated cluster accounted for 30.0% of MCseg cells (326 of 1,085), an increase of 2.7 percentage points. These percentages describe cluster assignment rather than a direct measurement of cell-recovery sensitivity. Scale bar: 50 µm.

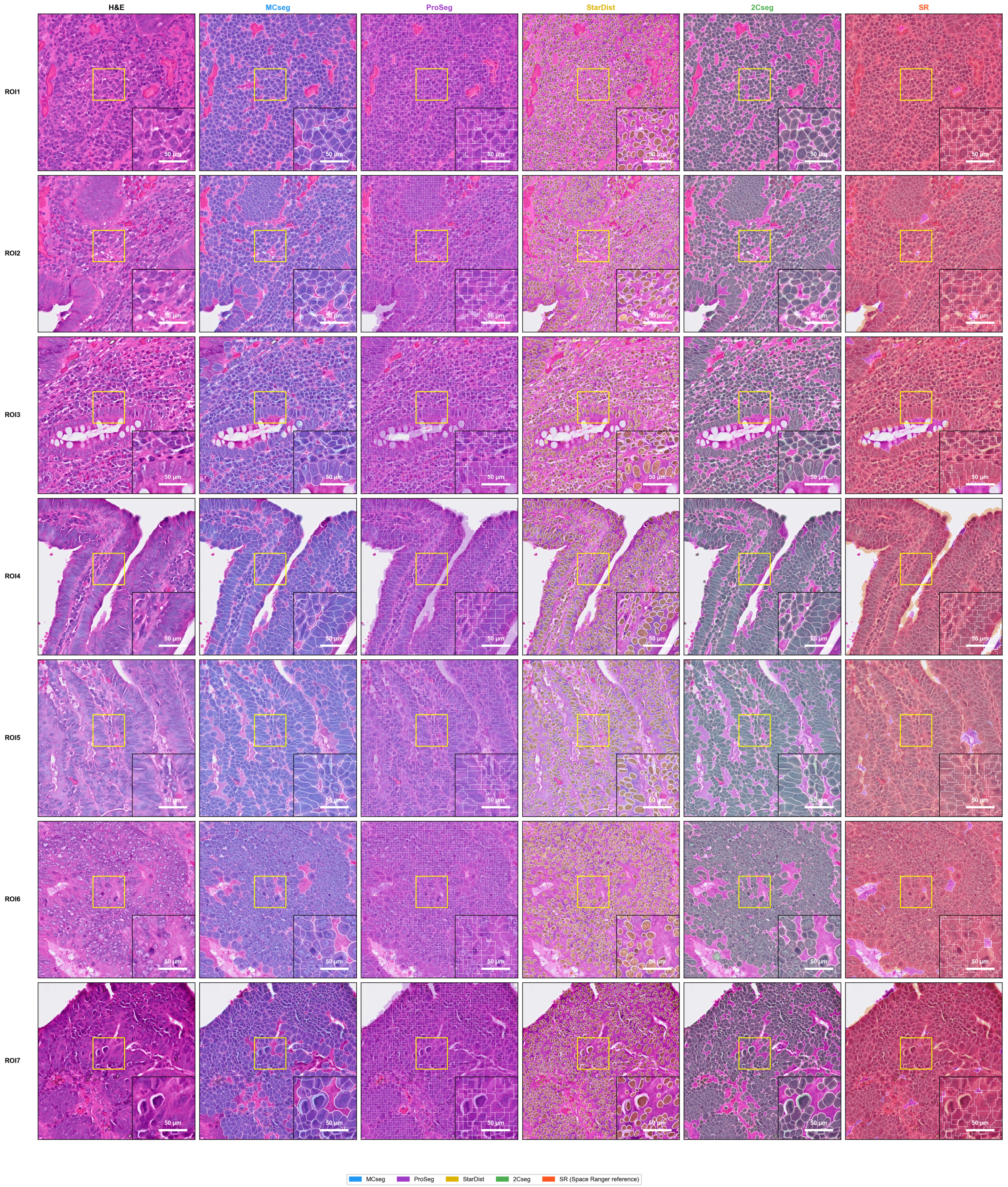


**Supplementary Figure S6.** Visual segmentation comparison across seven representative CRC ROIs. Columns show H&E, MCseg, Proseg, StarDist, 2Cseg, and Space Ranger (SR). Insets highlight representative local differences. MCseg generally maintained tighter cell-conforming boundaries than SR and left more inter-mask space in luminal or stromal regions. The qualitative comparison should be interpreted together with the transcript-derived metrics in Figure 3 and Supplementary Table S1. Scale bars: 50 μm.

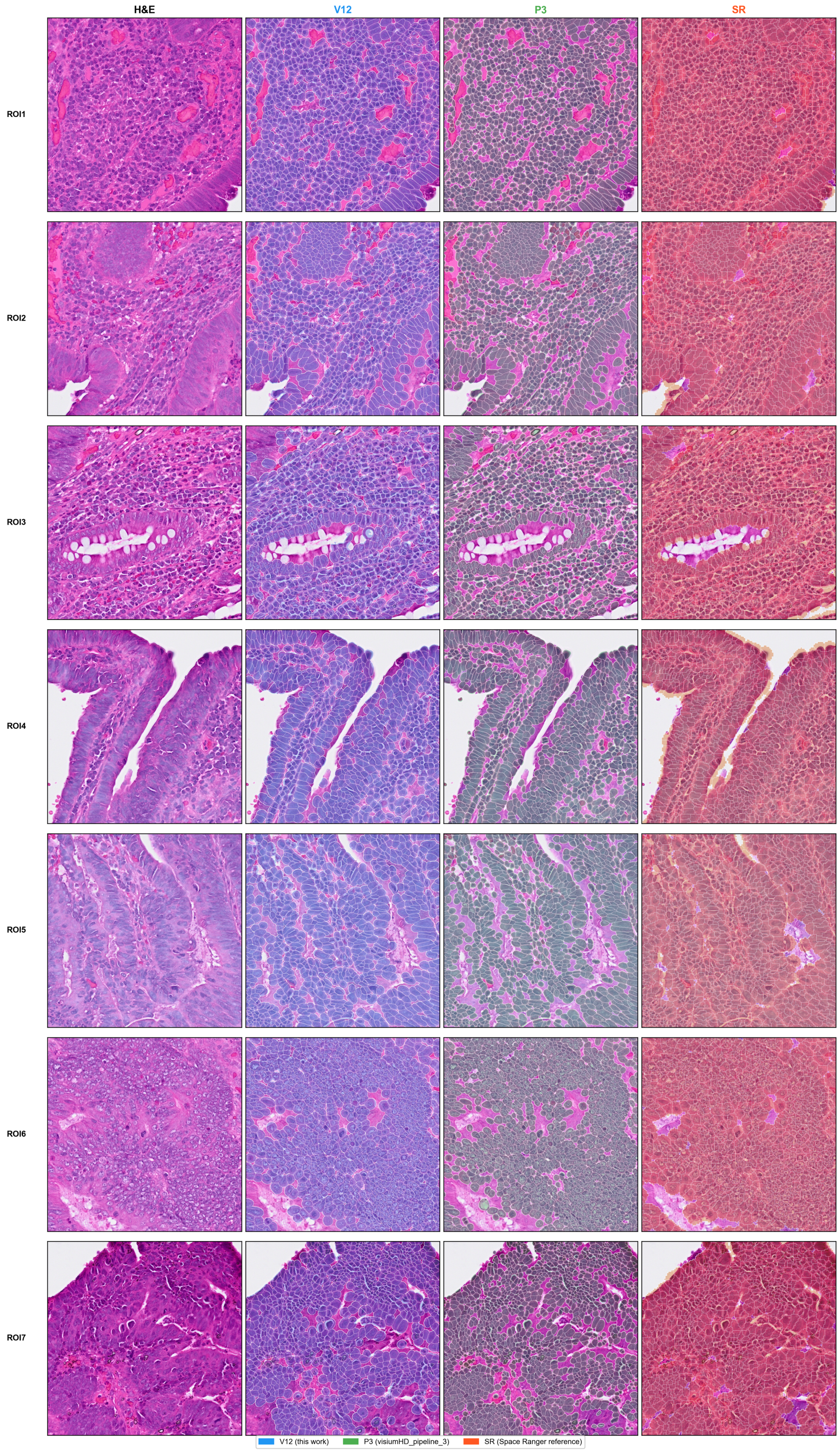


**Supplementary Figure S7.** Qualitative comparison of the final MCseg deployment architecture with an earlier development prototype and Space Ranger across seven representative CRC ROIs. Columns show H&E, final MCseg, the earlier prototype, and Space Ranger. The internal labels V12 and P3 in the original analysis outputs correspond to final MCseg and the earlier prototype, respectively. This figure illustrates the visual progression of the development workflow and is not a quantitative ablation experiment.

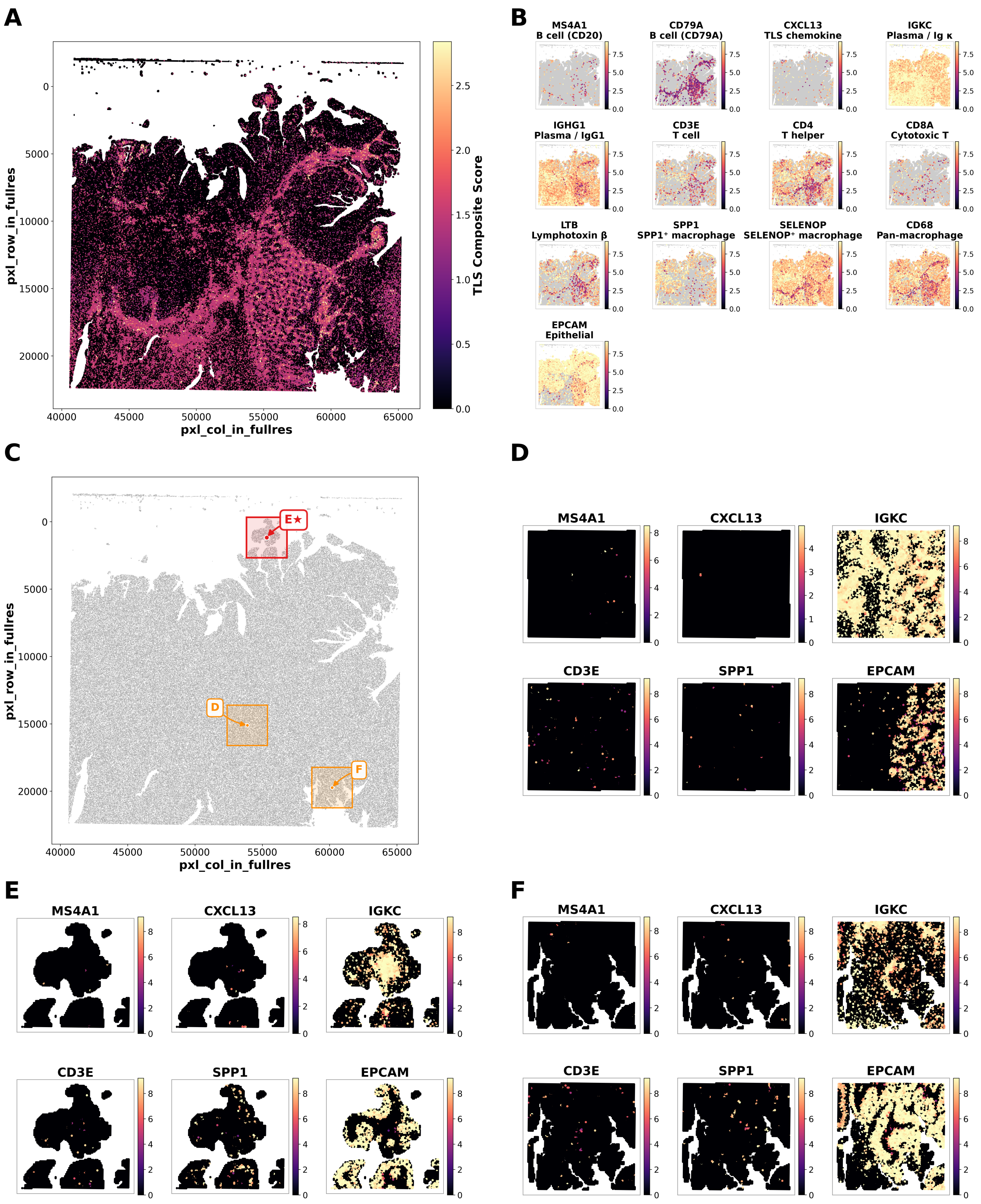


**Supplementary Figure S8.** TLS candidate discovery and selection on the CRC slide. (a) Full-slide composite score calculated from MS4A1, CD79A, CXCL13, IGKC, CD3E, and LTB across 8-µm bins. (b) Full-slide marker-gene panel used to interpret the composite signal. (c) Significant local Moran's I high–high hotspots and the three candidate regions taken forward for inspection. Candidate E was selected as ROI 15; candidate D was the largest hotspot but was not selected. (d–f) Zoomed marker maps for the three candidates, showing MS4A1, CXCL13, IGKC, CD3E, SPP1, and EPCAM. Local Moran's I used 8,999 permutations with $p < 0.01$.

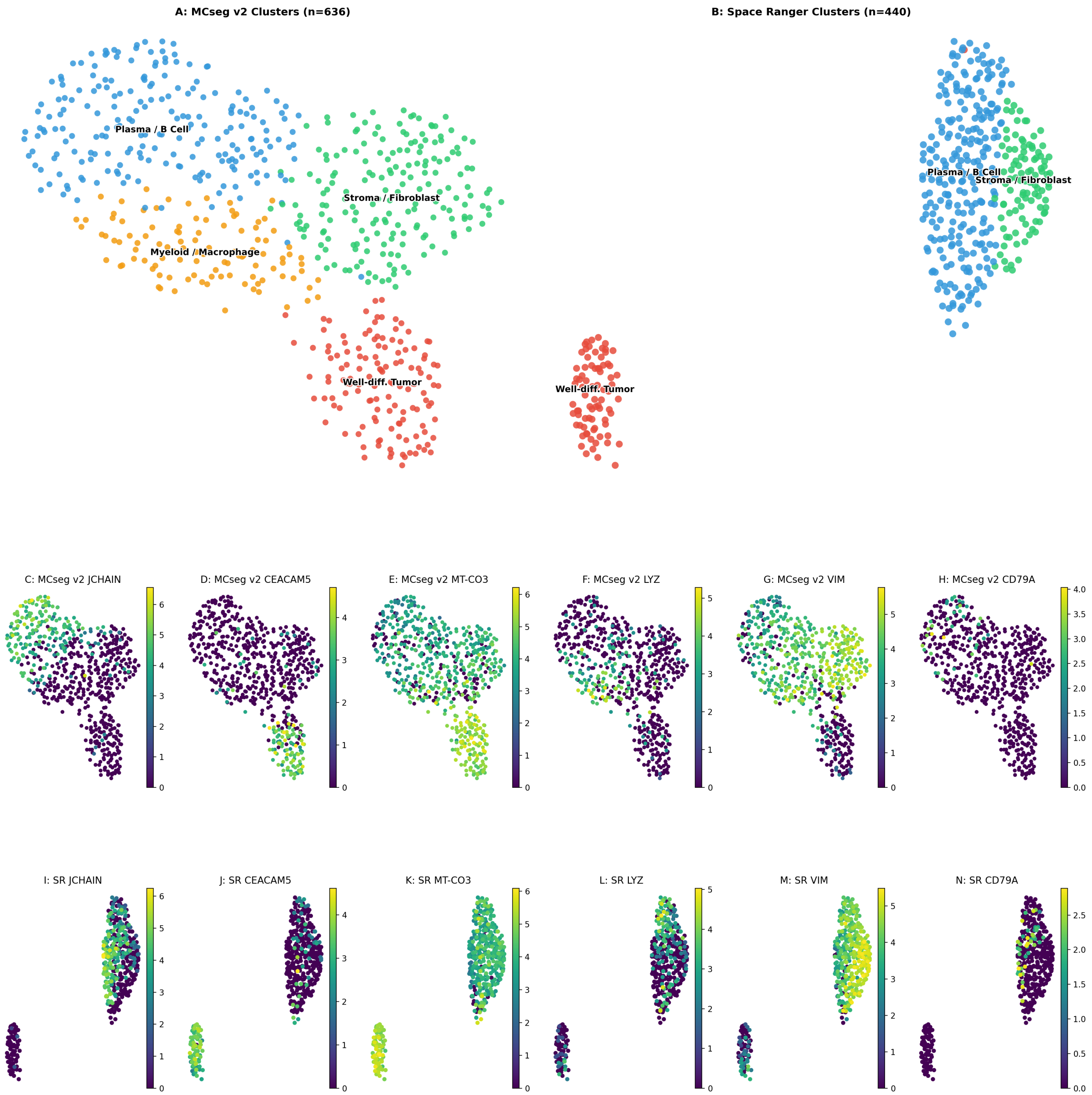


**Supplementary Figure S9.** UMAP analysis of CRC TLS ROI 15. (a) MCseg clusters (n = 636 cells; Leiden resolution 0.5), showing plasma/B-cell, stromal/fibroblast, myeloid/macrophage, and well-differentiated tumor populations. (b) Space Ranger clusters in the same region (n = 440 cells; Leiden resolution 0.5). (c–h) MCseg expression of JCHAIN, CEACAM5, MT-CO3, LYZ, VIM, and CD79A. (i–n) Expression of the same genes after Space Ranger segmentation.

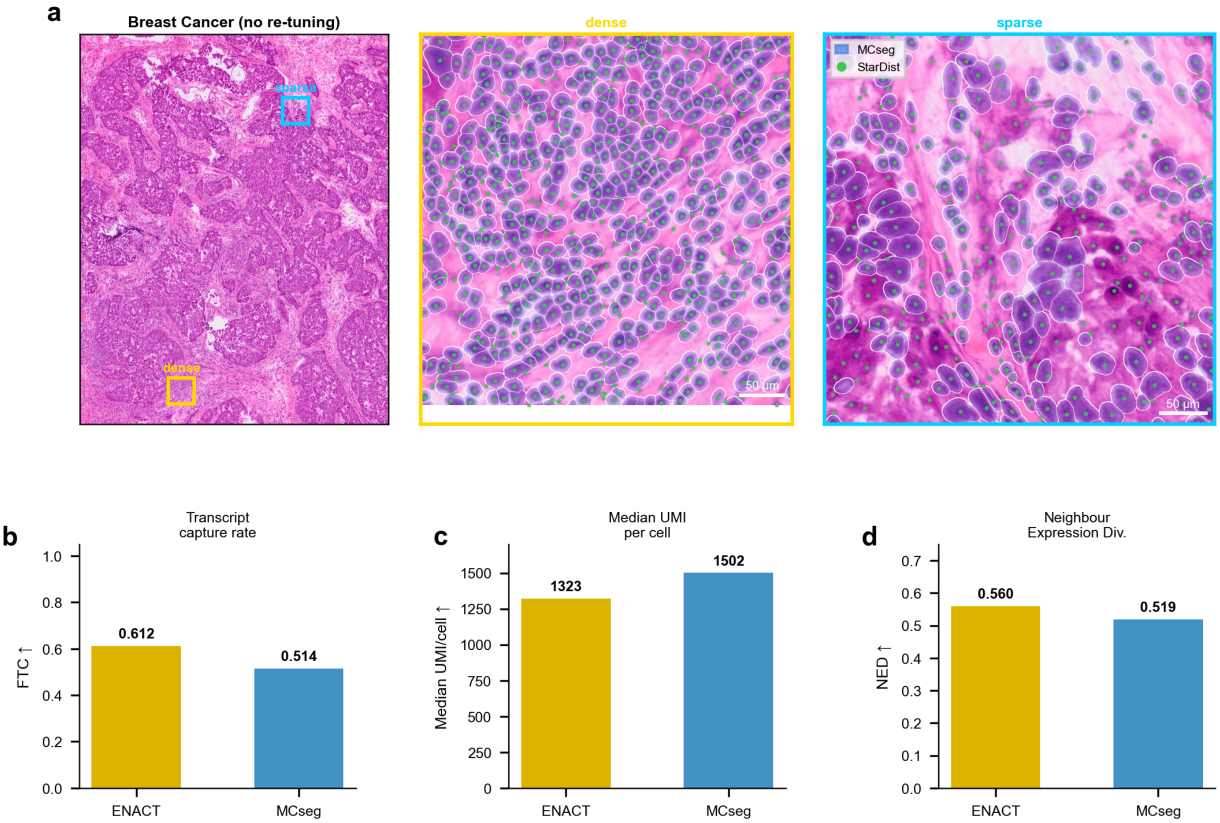


**Supplementary Figure S10.** Fixed-architecture application to a 10x Genomics Visium HD fresh-frozen breast cancer demonstration dataset. MCseg was applied without breast-specific architecture search. (a) Whole-crop H&E overview and representative dense and sparse regions. MCseg boundaries are shown in blue, and green dots indicate the StarDist centroids used by the ENACT comparator. (b) FTC: 0.514 for MCseg and 0.612 for ENACT. (c) Median UMI per cell: 1,502 for MCseg and 1,323 for ENACT. (d) NED: 0.519 for MCseg and 0.560 for ENACT. MCseg segmented 96,876 cells, whereas the ENACT StarDist-based comparator segmented 185,483 cells. Differences in per-cell UMI counts may reflect cell number, mask geometry, and transcript aggregation; MCseg does not use fractional boundary-bin sharing. These results provide a transfer test rather than evidence of superiority.

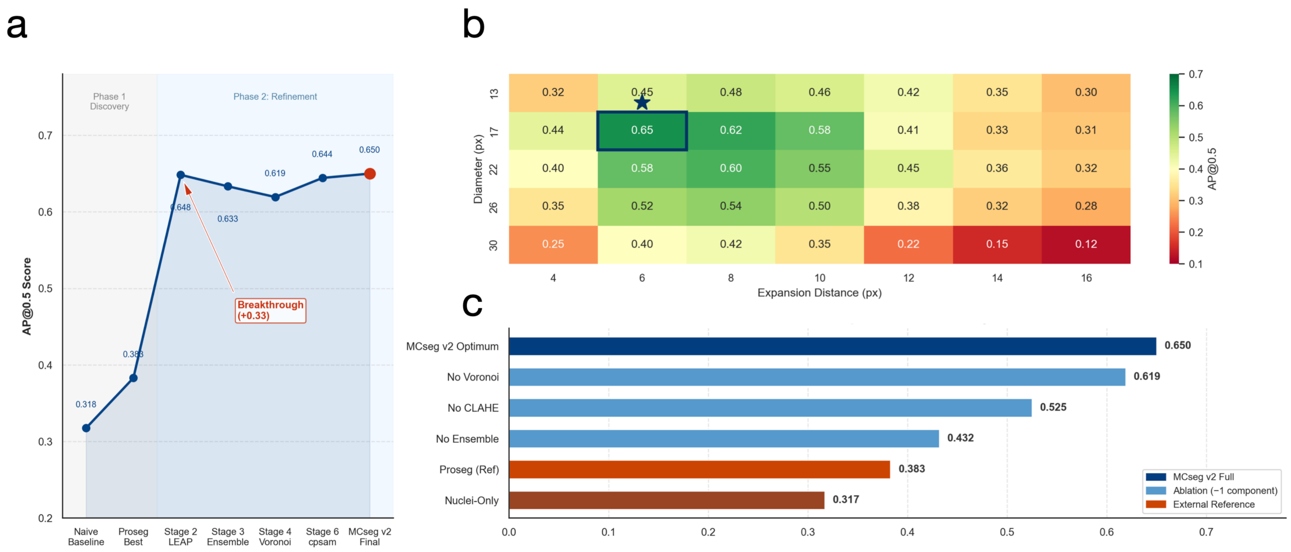


**Supplementary Figure S11.** AI-agent architecture search: performance trajectory, parameter sensitivity, and component contributions. (a) Evolutionary discovery path. AP@0.5 on the primary development patch across successive stages of the AI-agent-guided search, from the naive single-pass Cellpose baseline and the external Proseg comparator through intermediate architectures to the retained seven-pass configuration (AP@0.5 = 0.650). (b) Parameter sensitivity heatmap. AP@0.5 as a function of Cellpose diameter and Voronoi expansion distance sampled during the search; the retained configuration falls within the highest-scoring region of the sampled space. (c) Component ablation study. AP@0.5 for the full retained architecture compared with variants omitting individual components (no Voronoi expansion, no CLAHE preprocessing, no multi-pass mask integration) and with external references (Proseg, nuclei-only segmentation). These scores describe the development patch used during architecture search and are distinct from the six-ROI fixed-parameter deployment benchmark (PQ) reported in the main text (Results 3.2).
